# MiRNA-10b marks aggressive squamous cell carcinomas, and confers a cancer stem cell-like phenotype

**DOI:** 10.1101/2020.02.05.934109

**Authors:** Monika Wimmer, Roland Zauner, Michael Ablinger, Josefina Piñón-Hofbauer, Christina Guttmann-Gruber, Manuela Reisenberger, Thomas Lettner, Norbert Niklas, Johannes Proell, Mila Sajinovic, Paul De Souza, Stefan Hainzl, Thomas Kocher, Eva M. Murauer, Johann W. Bauer, Dirk Strunk, Julia Reichelt, Albert S. Mellick, Verena Wally

**Author notes:** MW and RZ equally contributed to the work. **Corresponding author:** Verena Wally.

## Abstract

**Background:** Cutaneous squamous cell carcinomas (cSCC) are the primary cause of premature deaths in patients suffering from the rare skin-fragility disorder recessive dystrophic epidermolysis bullosa, which is in marked contrast to the rarely metastasizing nature of these carcinomas in the general population. This remarkable difference is attributed to the frequent development of chronic wounds caused by an impaired skin integrity. However, the specific molecular and cellular changes to malignancy, and whether there are common players in different types of aggressive cSCCs, remain relatively undefined.

**Methods:** MiRNA expression profiling was performed across various cell types isolated from skin and cSCCs. Microarray results were confirmed by qPCR and by an optimized *in situ* hybridization protocol. Functional impact of overexpression of a dysregulated miRNA was assessed in migration and 3D spheroid assays. Sample-matched transcriptome data was generated to support the identification of disease relevant miRNA targets.

**Results:** Several miRNAs were identified as dysregulated in cSCCs as compared to controls. These included the metastasis-linked miR-10b, which was significantly upregulated in primary cell cultures and in archival biopsies. At the functional level, overexpression of miR-10b conferred the stem cell-characteristic of 3D-spheroid formation capacity to keratinocytes, and impaired their mobility. Analysis of miR-10b downstream effects identified a novel putative target of miR-10b, the actin- and tubulin cytoskeleton-associated protein DIAPH2.

**Conclusion:** The discovery that miR-10b confers an aspect of cancer stemness – that of enhanced tumor cell adhesion, known to facilitate metastatic colonization - provides an important avenue for future development of novel therapies targeting this metastasis-linked miRNA.

## Background

While the incidence of cutaneous squamous cell carcinomas (cSCC) in Western industrialized societies has increased in recent years, it has remained relatively easy to treat, especially at early stages of the disease (1). However, in some cases cSCCs metastasize, worsening the prognosis of patients. A special case of cSCC occurs in patients suffering from the rare genodermatosis recessive dystrophic epidermolysis bullosa (RDEB). These patients are at high risk of developing a particularly aggressive type of cSCC with a high metastatic potential that is linked to changes in the extracellular matrix (ECM), caused by loss-of-function mutations in the *COL7A1* gene (2). A lack of functional type-VII collagen (C7) at the dermal-epidermal junction (DEJ) sensitizes skin to blistering, and erosions, upon minor physical stress or trauma (3). Patients with RDEB present with congenital generalized blistering and a number of severe secondary manifestations. cSCCs arise in nearly all RDEB patients by the age of 45, and associated metastatic disease is the primary cause of premature deaths in RDEB patients (4). Several mechanisms are thought to contribute to the aggressive and rapidly progressing nature of RDEB-SCCs. In general, the skin’s constant need to repair itself, coupled with the stalled inflammatory processes, and aberrant TGF-ß signaling associated with microbial challenge (5-9), are considered major risk factors. To which extent these inflammatory changes are linked to the particularly aggressive form of cSCC associated with RDEB, and if these tumors have characteristics in common with cSCCs that present with an aggressive behaviour in otherwise healthy people, remains unknown.

We focused on post-transcriptional regulatory processes in aggressive cSCCs, in particular on micro-RNAs (miRNAs). miRNAs are short (20 – 25 nucleotide) RNA molecules, which are key regulators of normal cell functions. In a healthy system, miRNAs are predicted to mediate the post-transcriptional control of up to 60% of all expressed genes (10). Their dysregulation is associated with several pathologic states, including cancer, heart disease, and obesity, and they are attributed a promising potential for therapeutic developments (11,12). In recent years, both, oncogenic miRNAs (onco-miRs) and tumor suppressive miRNAs, have been identified as playing important roles in cancer progression. In addition, a class of miRNAs have been shown to have specific pro-metastatic properties. A key metasta-miR, miR-10b, has been attributed with tumor promoting properties, as well as the growth of metastatic foci in breast cancer in various landmark studies (13-15). *MiR-10b* is encoded by a highly conserved genomic region, which is located near the homeobox D (HOXD) cluster on chromosome 2 (16). It has been linked to a range of functions, including regulation of angiogenesis and promotion of cell invasion (17-20). In addition, increased serum levels of miR-10b are associated with poor prognosis in melanoma and breast cancer (21,22), and intravenous injections of miR-10b inhibitors in tumor-bearing mice dramatically reduced breast cancer cell metastasis (13,15,23). In addition, a meta-analysis of miR-10b levels and clinical outcomes in various cancers, demonstrated that overexpression was associated with poor overall survival, indicating that miR-10b might be a promising prognostic biomarker (24).

In this study, we substantially add to our understanding of the role of miR-10b, by reporting for the first time on the dysregulation of this miRNA in aggressive cSCC. Both, RDEB- and otherwise healthy donor-related cSCCs (HC-cSCCs) were classified and included in this study according to their potential to metastasize. We further show that miR-10b expression is linked to a cancer stem cell-like phenotype in a 3D organotypic model. Taken as a whole, this work provides a new explanation of maligancy in cSCCs, and a novel target for further development of markers and therapies to treat cSCCs.

## Methods

### Patient samples and cell lines

For this study, primary cells were used for microarray experiments, as well as for validations. For miR-10b overexpression, we used HPV16 E6/E7 immortalized keratinocytes. All cells were cultured in defined, serum-free CnT-Prime Epithelial Culture Medium (CELLnTEC) at 37°C / 5% CO_2_ in a humidified incubator. For more detailed information regarding origin of the cell lines and donor description see Supplementary Information and Supplementary Table S1.

### Microarrays

For total RNA and miRNA isolation from cell cultures, we used miRNeasy Mini Kit (Qiagen, 217004), according to manufacturer’s protocol. For transcriptome analysis, an Affymetrix Clariom™D (Thermo Fisher, 902922) was applied and an Affymetrix GeneChip™ miRNA 4.1 platform was used for miRNA analysis (Thermo Fisher, 902409). Arrays were performed by the service provider “Core Facility Genomics at the Medical University Vienna” in accordance with manufacturer’s instructions. RNA quality was confirmed on a Bioanalyzer prior to hybridization onto a respective array. Quality assessment of microarray data was conducted via “Transcriptome Analysis Console” (Applied Biosystems v4.0.0.25).

### Bioinformatic data processing and statistical analysis

All data processing and analysis was documented and performed in statistical software R (v3.5.1). Scripts are available upon request. For detailed statistical analysis refer to Supplementary Information.

For statistical analysis Student’s t-test was performed using the GraphPad Prism (v 5.03) or Excel software, and error bars represent standard error of mean (SEM), unless mentioned otherwise. Details regarding number of replicates can be found in figure legends.

### TaqMan qPCR miRNA assays

For primary (pri)-miRNA assays cDNA was synthesized from 1 µg total RNA. For reverse transcription, the High Capacity RNA-to-cDNA kit (Applied Biosystems, 4387406) was used according to the manufacturer’s protocol. TaqMan pri-miRNA assays (Thermo Fisher Scientific, 4427012, Hs03302884_pri) including the TaqMan Gene Expression Master Mix, were used. RNaseP (Thermo Fisher Scientific, 4401631) served as endogenous control.

For mature miRNA validations we used TaqMan Advanced miRNA assays (Thermo Fisher Scientific, A25576), 2x Fast Advanced Master Mix, and respectively diluted miRNA templates. miR-320a-3p (Thermo Fisher Scientific, 478594_mir) was used as endogenous control. All reactions were performed in technical triplicates, and repeated as independently performed experiments for at least three times.

### Fluorescence-based in situ hybridization of miR-10b on cultured cells and formalin-fixed paraffin embedded (FFPE) tissue sections

An optimized *in situ* hybridization protocol was applied, based on the method described by Gasch *et al*. (13) and the manufacturer’s recommendations (Qiagen, 339450). In brief, 8 µm FFPE tissue sections were deparaffinized in xylene, washed and rehydrated in serial dilution of ethanol / dH_2_O. Antigen retrieval was performed in citrate buffer pH 6.0 at 100 °C.

Cells were cultured in 8-well glass chamber slides (Millipore) and fixed with 4 % PFA in PBS at room temperature, and permeabilized by incubation in 1 % Triton-X100 in PBS. After re-fixation in 4 % PFA in PBS, cells were washed twice in PBS and incubated for 20 min at 54 °C (60 min at 52 °C for tissue sections) in mercury locked-nucleid acid (LNA) miRNA ISH Buffer 80 nM hybridiziation buffer (100 nM for tissue section) (Qiagen, 339450). Digoxigenin labeled miRCURY LNA probes (HSA-MIR-10B-5P, SCRAMBLE-MIR, miRCURY LNA U6, Qiagen) were diluted in hybridiziation buffer after a denaturation step at 90 °C and incubated on cells for 30 min (60 min tissue sections) at 54 °C (52 °C tissue sections). After probe hybridization increasingly stringent washing steps were applied in saline-sodium citrate (SSC) buffer (Sigma, SSC 20x buffer concentrate S6639-1L) at various concentrations and temperatures. Tris based (100 mM) buffer was supplemented with 3 % (v/v) FCS, 1 % (w/v) hyridization blocking reagent (Roche, 339450), 0.3 % (v/v) Triton-X100 and NaCl (150 mM). Anti-DIG Fabs (Roche, 11207741910), Alexa Fluor® 647 Mouse Anti-Human Cytokeratin 14, 15, 16 and 19 Clone KA4 (Supplementary Table S2) and DAPI were used for visualization and co-staining of cells and tissue section, respectively. Slides were mounted using Prolong Gold Antifade mounting medium (Sigma, P36930).

Imaging was performed on a LSM800 Airyscan confocal microscope (Zeiss). Analysis was conducted in ImageJ (v1.52i) and CellProfiler (v3.0.0) with all steps documented in scripts and workflow files, available upon request. Cellular boundaries were detected based on respective fluorescence signals and signal intensities were integrated at single cell resolution. To confine the analysis to cytoplasmic signals, nuclei were demarked via DAPI staining and co-localizing fluorescence signals were subtracted from total fluorescence. Downstream statistical analysis was performed in statistical software R (v3.5.1).

### Histological analysis of tissue sections and immunofluorescence microscopy

cSCC samples from patients undergoing tumor resection were FFPE processed (25). Hematoxilin & eosin (H&E) stainings were performed for histological assessment by a dermato-histopathologist (Department of Dermatology, University Hospital Salzburg).

For immunohistochemistry (IHC), sections were incubated at 58°C for 1 hour, and subsequently deparaffinized three times in Roti-Histol (Roth, 6640) at room temperature (RT). Next, sections were washed in decreasing alcohol concentrations (3 × 100 % EtOH, 2 × 90 % isopropanol, 1 × 70 % isopropanol, 1 × dH_2_O). Heat-induced antigen retrieval was performed in citrate buffer pH 6.0. Upon blocking with 5% bovine serum albumin (BSA, Sigma, A3294) in PBS, first antibodies were diluted in 2 % BSA/PBS. For all antibodies, a respective second-step control was included. Secondary antibodies were used at a 1:500 dilution (Supplementary Table S2). DAPI was used for nuclear staining (Sigma Aldrich, D9542). The same protocol was used for cultured cells. There, about 3 × 10^4^ cells were seeded into 1 cm^2^ chamber slides. After 24 hours (hrs) cells were fixed for 10 min in 4 % formaldehyde in PBS, washed once with PBS and blocked at room temperature for one hour with 2 % BSA in PBS. Imaging was performed on a Zeiss LSM710 confocal microscope.

### Cloning of miR-10b and stable transduction

For stable expression of miR-10b in E6/E7 immortalized HC-KCs and RDEB-KCs, the human *MIR10B* gene was cloned into the pMX-IRES-Blasticidin vector (Cell Biolabs Inc., RTV-016), downstream of the constitutive Pol-III U6 promoter. Primer sequences are given in Supplementary Table S3. All constructs were analyzed using Sanger sequencing before viral packaging.

Viral particle production using pMX_U6_miR10b was done as described previously (26). Expression and maturation of miR-10b was confirmed by TaqMan-PCR (Supplementary Fig. S1C-E).

### CRISPR-mediated knock-out of MIR10B

Two single guide (g)RNAs, flanking the *MIR10B* stem loop region on chromosome 2, were rationally designed and selected to specifically knock-out *MIR10B*. Single gRNAs were synthesized and purified using the Precision gRNA Synthesis Kit according to the manufacturer’s instructions (Thermo Fisher Scientific, A29377). Two primer sets were used for gRNA synthesis (Supplementary Table S3). Recombinant spCas9 was purchased from Polyplus-transfection (Polyplus-transfection). Ribonucleoproteins (RNPs) were complexed in a 4:1 ratio (3 µg spCas9 and 750 ng sgRNAs; 375 ng each sgRNA) for 10 min at room temperature and delivered into RDEB-SCC1 keratinocytes by nucleofection using the Neon Transfection System 10 μL Kit (Thermo Fisher Scientific, MPK10025). SCC cells were trypsinized and washed with PBS, and 3×10^5^ keratinocytes were resuspended in 12 μL Resuspension Buffer R per reaction. RNP complexes were added to each sample and electroporated into SCC cells (∼2.5×10^5^ cells) under the following conditions: 1400 V, 20 ms, 2 pulses. After electroporation, cells were seeded into 6-well plates containing pre-warmed antibiotic-free CnT-Prime Epithelium Culture Medium (CELLnTEC).

### Migration assay

Cell motility assays were performed using 2-well silicone inserts with a defined cell-free gap, suitable for wound healing / migration assays (IBIDI, 80241). 7×10^4^ cells in 70 µL CnT-Prime medium were seeded per well. 24 hrs post seeding silicone inserts were removed and migration of cells was monitored at different time points by measuring cell confluence in a pre-defined, constant area including the gap using a Spark^®^ 10M multimode microplate reader imaging module (Tecan). Gap-area was calculated using ImageJ (v1.52i). In parallel to every migration assay, proliferation of each cell type was measured in 24-well plates, to exclude varying proliferation rates as confounding factor.

### Generation of 3D tumor spheroids

3D spheroids were grown as floating spheres in AggreWell™ 400Ex Plates (Stem-cell Technologies, 34425) according to the manufacturer’s recommendations. 2.35×10^5^ cells / well were seeded in CnT-prime medium supplemented with 1.2 mM CaCl_2_. Spheroids were grown for 96 hrs. Aggregates were transferred into a 24-well plate for imaging on a Spark^®^10M multimode microplate-reader with integrated imaging module (Tecan). For live/dead stainings, aggregates were purified using Corning^®^ Costar^®^ Spin-X^®^ Centrifuge tube filters (0.22 µm) (Corning, CLS8161-100EA). Viability was tested via live/dead viability/cytotoxicity kit (Thermo Fisher, L3224). Imaging was performed on a Zeiss LSM710 confocal microscope. TECAN whole well images were processed in ImageJ (v1.52i) software. All image processing was documented in ImageJ macros and downstream statistical analysis in R (v3.5.1) scripts, which are available upon request.

For outgrowth experiments, spheroids were transferred to standard 24-well plates and incubated at 37°C in a humidified incubator using CnT-Prime Epithelium Culture Medium. Cellular outgrowth was imaged 24 hrs post transfer.

### Semi-quantitative real-time PCR

SqRT-PCR was performed using the GoTaq® qPCR Master Mix (Promega, TM318), according to the manufacturer’s protocol. Glycerinaldehyd-3-phosphat-dehydrogenase (*GAPDH*) and tubulin alpha 1 (*TUBA1*) were used as reference genes. For primers see Supplementary Table S3. Data collection was performed on a CFX96 (BioRad) instrument. Relative target gene expression quantities and significance levels (unpaired two-sided t-test) were calculated using the ΔΔCq method. All reactions were performed in technical duplicates, and performed in at least three independent experiments.

### Western blot analysis

Protein extracts were generated from ∼ 5×10^5^ cells at a confluency of 80%, dissolved and homogenized in RIPA buffer (Santa Cruz, sc-24948) supplemented with 5% ß-Mercaptoethanol (ß-ME). Proteins were separated on a 4-15% bis-tris gel (Invitrogen, Nupage, NP0323) in MOPS buffer and blotted onto a 0.45 µm nitrocellulose membrane (Amersham Biosciences, Hybond-ECL, Merck, GERPN303D). Ponceau Red staining was performed for total protein analysis of ChemiDoc (BioRad) images in ImageLab software (BioRad, v5.2.1). Membranes were blocked using 5% milk powder in TBS / 0.1% Tween20 (Merck, 655205-250ML). All antibodies are listed in Supplementary Table S2. Membranes were washed five times using TBS / 0.1% Tween-20. HRP activity was visualized using the Amersham ECL Select Western blot detection reagent (Amersham Biosciences, RPN2235), and the ChemiDoc system (BioRad). Specificity of the DIAPH2 antibody was confirmed in DIAPH2 knock-out cells. Dosimetric semi-quantitative analysis was performed in either ImageLab (BioRad, v5.2.1) or ImageJ (v 1.52i).

### MiR-10b mimic transfections

Primary healthy control keratinocytes (HC-KC) were transiently transfected using Xfect transfection reagent (TaKaRa Bio, 631450/631317) with 50 nM miR-10b-5p mimic / ∼10^5^ cells (Thermo Fisher, miRVana, 4464066) or scrambled control (Merck, HMC0002) according to manufacturer’s protocol. Cells were lysed 72 hrs post-transfection in RIPA buffer (Santa Cruz, sc-24948) supplemented with 5% ß-ME for subsequent Western blot analysis.

## Results

### MiRNome profiling identifies upregulation of miR-10b in RDEB-SCC

In order to identify differences in the miRNA expression profile between cultured primary keratinocytes (KC) and cSCC cells (Supplementary Information and Supplementary Table S1), we conducted an Affymetrix GeneChip™ miRNA 4.1 expression analysis. Out of 2,578 analyzed mature miRNAs unique to human, 50 miRNAs were found to be significantly (p-value ≤ 0.05 and false discovery rate (FDR) ≤ 0.2) ≥ 2-fold up- or down-regulated in RDEB-SCCs (n = 4) compared to RDEB-KC (n = 6), and 56 in cSCCs from otherwise healthy donors (HC-cSCCs) compared to healthy control (HC)-KCs (n = 5, Supplementary Fig. S1A, Supplementary Tables S4 and S5). When analyzing miRTarbase v6.1 predicted targets of the deregulated (≥ 2-fold, p-value ≤ 0.05 and FDR ≤ 0.2) miRNome, various cancer-related Kyoto Encyclopedia of Genes and Genomes (KEGG) pathways, *e.g*. signaling pathways regulating pluripotency of stem cells, appeared significantly enriched (Supplementary Fig. S1B).

To assess the ability of the miRNA expression profiles to distinguish cSCCs from other experimental groups we performed principal component analysis (PCA). Annotation of the samples within the distinct clusters resulted in a clear separation of experimental groups (Figure 1A). Even though the miR-10 family was found to be overall upregulated in cSCCs, in-depth analysis of major contributors driving the unsupervised cluster separation highlighted miR-10b, which was 2.2-fold (p < 0.05) upregulated in RDEB-SCCs and 2.2-fold (p < 0.05) in HC-cSCCs, respectively, compared to their non-malignant controls (Figure 1B,C). As opposed to miR-10b, miR-10a was only found to be slightly upregulated by qPCR. As miR-10a appeared to be the most deregulated miRNA in RDEB-SCCs in microarray analysis but not in subsequent qPCR, we assume that high microarray scores most likely derived from a certain hybridization error rate (miRs-10a and −10b differ in only one nucleotide) (Figure 1E, Supplementary Fig. S1C-E). In addition, increased miR-10b (9-fold), but not miR-10a, levels in RDEB-SCC were found in previously generated RNA sequencing (RNA-seq) data, where immortalized HC-KC lines were used as controls (Supplementary Fig. S1F,G). Thus, we focused on miR-10b, and dropped miR-10a from further experiments. Predicted miR-10b targets were further tested for gene set enrichment in cancer hallmarks (Molecular Signature Database v6.2), and showed significant association with metastatic processes like epithelial to mesenchymal transition (EMT) (Figure 1D).

**Figure 1:**
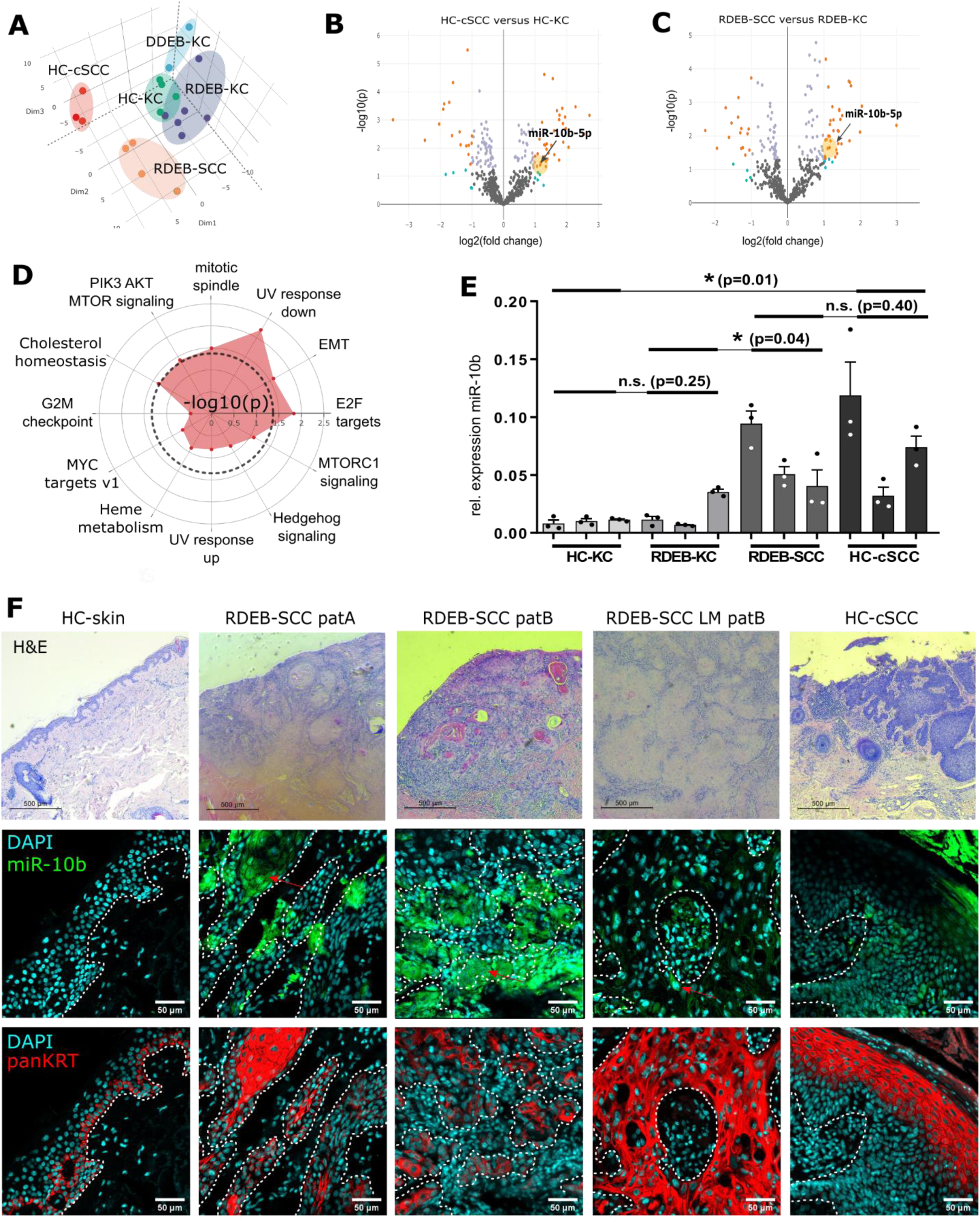
miR-10b is upregulated in RDEB-SCC. (A) Unsupervised clustering via PCA on miRNA expression levels clearly separated RDEB-SCC (n = 4) and HC-cSCC (n = 3) samples from RDEB-KC (n = 6) and HC-HC (n = 4). Target analysis of miR-10b, which was significantly (p ≤ 0.05) up-regulated (≥2-fold) in both, (B) HC-cSCCs and (C) RDEB-SCCs, indicated an enrichment of pathways related to aggressive tumor phenotypes like epithelial-to-mesenchymal transition (EMT) (D). (E) TaqMan qPCR (n=3 ind. repl., mean ± SEM, *p < 0.05, n.s. non-significant, unpaired t-test), as well as (F) *in-situ* hybridization (FISH) with DIG labelled miR-10b probes (green) on FFPE tissues sections of confirmed (H&E staining, top row) carcinomas also showed miR-10b dysregulation. (red: tumor boundaries marked by pan-keratin staining) Scale bars: 100μm (white), 500μm (black).

In summary, the results of a microarray-based miRNA expression profiling revealed a deregulated miRNome able to distinguish cSCC from keratinocytes. In particular, miR-10b was significantly upregulated in cSCC.

### Expression of miR-10b in cSCC tissues

In order to localize and further substantiate upregulation of miR-10b expression in tissue, we optimized a combined IHC-ISH protocol using LNA probes. When tissue specific expression was examined in FFPE-sections of archival skin and tumor biopsies, miR-10b was found to be upregulated in cSCC, particularly in RDEB-SCCs. Expression was predominantly co-localized to keratin-positive cells (Figure 1F, Supplementary Fig. S2A-C), tumor vasculature, and lymphocytes (Supplementary Fig. S2D). Notably, both vasculature and hemopoietic cells have previously been shown to express miR-10b in a tumor context (15, 18, 27-29). Further analysis of RDEB-tissues showed a strong expression of transcription factor TWIST1, a known upstream driver of miR-10b-mediated tumor malignancy. In addition, TWIST1 was also found to be upregulated in cultured SCC cells in IF and Western blot analysis (Supplementary Fig. S2E,F) (14, 30). Single cells in an RDEB-SCC lymphnode metastasis (RDEB-SCC_LM) expressed high levels of miR-10b. Less pronounced miR-10b abundance was observed in HC-cSCC (Figure 1F).

To investigate whether miR-10b expression was correlated to differentiation, cells were incubated in the presence of calcium and serum in order to induce differentiation, and respective marker gene expression was analyzed in parallel to miR-10b expression levels. However, no overall correlation between differentiation and miR-10b expression was observed (data not shown).

Taken together, results of the IHC-ISH support a potential link between miR-10b expression levels and aggressive RDEB-SCC.

### MiR-10b confers anchorage-independent aggregation capabilities to keratinocytes and attenuates mobility

Metastasis requires the dissemination and successful establishment of clones with tumor-initiating potential at distant niches (31). In order to investigate the biological role of miR-10b, we analyzed its expression levels in experimental cell lines using IHC-ISH, in parallel to 3D tumor spheroid formation assays. As the cytoplasm is considered as the site of miRNA maturation and action, fluorescence intensity, excluding the nuclear region, was assessed at a single cell resolution. MiR-10b was confirmed to be highly expressed in RDEB-SCC cell lines using ISH probes specific for mature miR-10b, and in two out of three HC-cSCC cultures, as compared to primary HC-KCs (Figure 2A,B, Supplementary Fig. S3,4). Notably, we observed that miR-10b levels in cSCCs varied between individual cell lines, however, in correspondance to phenotypic features linked to increased malignancy, like an EMT-associated spindle-like phenotype.

**Figure 2:**
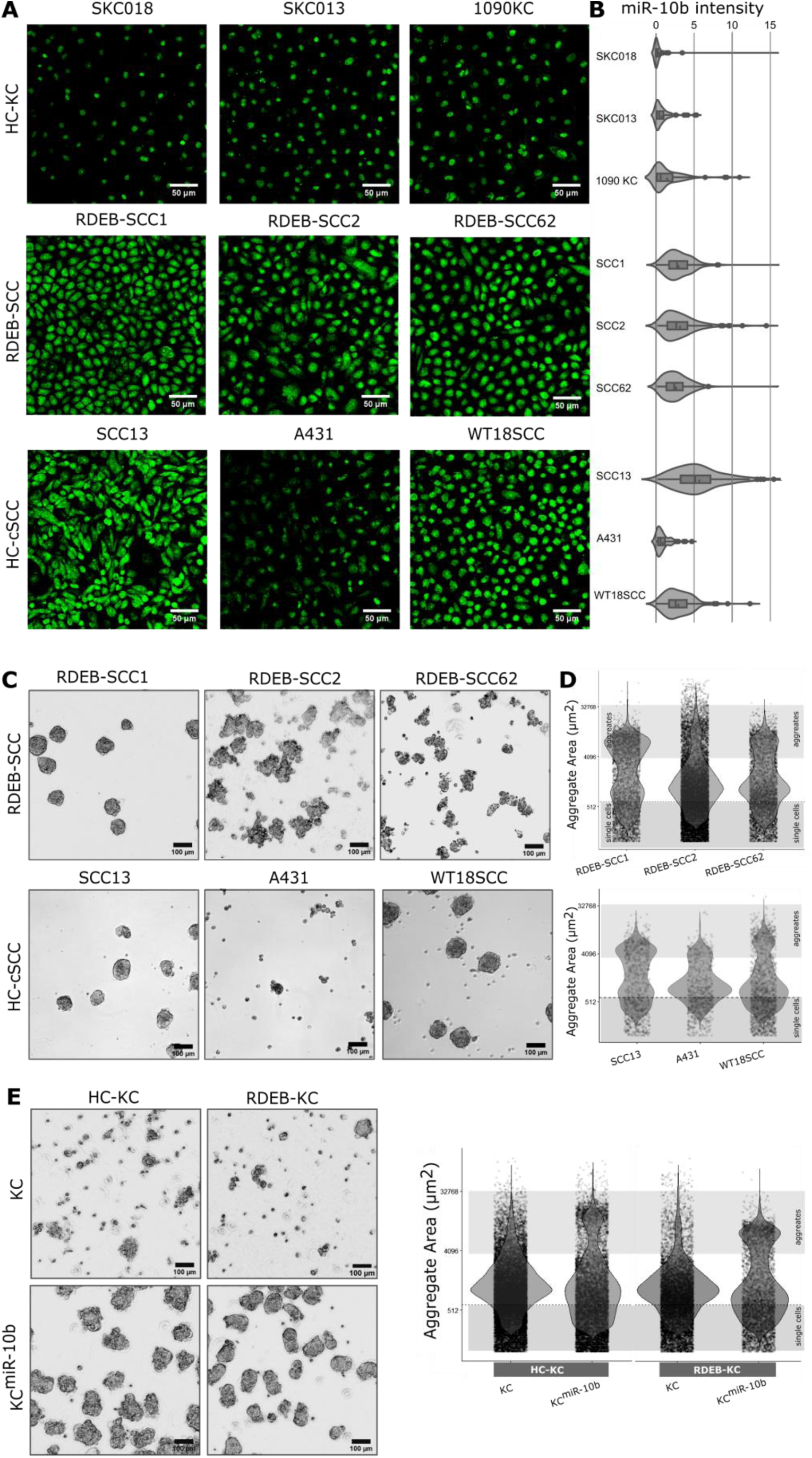
Stable aggregate formation is associated with miR-10b expression. (A) DIG-labelled miR-10b-5p specific LNA probes (green) were used in an *in-situ* hybridization protocol to quantify relative expression of miR-10b in cultured cells by FITC labeled anti-DIG-Fab immunofluorescence imaging. All three tested RDEB-SCCs and two out of three HC-cSCC cell lines demonstrated increased miR-10b signals compared to primary HC-KCs. (B) To quantify the miR-10b expression at single cell resolution, cytoplasmic FITC signals of cells were cumulated and plotted as relative fluorescence intensity per single cell on the x-axis. Graphic shows distribution density function of miR-10b intensity of all analyzed single cells per sample. Respective negative-(scrambled) and positive-(U6) probes were used as controls (Supplementary Fig. S4). (C) All RDEB-SCC lines and two out of three HC-cSCC lines demonstrated stable aggregate formation 96 hrs post seeding (representative images of aggregates). (D) Cross section of objects evaluated after transferring aggregates (n∼7·10^3^) into cell culture plates. Each dot represents one object. (E) In addition, overexpression of miR-10b in KCs conferred spheroid formation capacities. Scale bars: 50μm (black), 100μm (white).

Next, we performed aggregate formation assays to explore the capacity of our cell lines to form anchorage independent spheroids (32). All three RDEB-SCCs, and two out of three HC-cSCCs formed stable aggregates (Figure 2C,D). Notably, KCs transduced with, and stably expressing miR-10b (KC^miR-10b^) as surrogate for downstream functional experiments, phenocopied RDEB-SCCs in the formation of stable spheroids of 6.9×10^3^ ± 1.1×10^3^ µm^2^ (≙ 90 µm diameter, RDEB-KC, Figure 2E). Expression and maturation of miR-10b, driven by the constitutive Pol-III U6 promoter, was confirmed by qPCR (Supplementary Fig. S5A). In addition, we found that KC^miR-10b^ were significantly smaller than their parental cells (p-value^RDEB-KC^ = 0.002; p-value^HC-KC^ < 0.001; Supplementary Fig. S6A). Like in RDEB-SCC derived aggregates, viable cells were found predominantly in the outer spheroid layers of KC^miR-10b^ spheres (Supplementary Fig. S5I). Wild type KCs formed fewer and rather loose aggregates, which did not withstand mild pipetting.

In order to further substantiate the link between miR-10b overexpression and enhanced spheroid formation capacities, we next knocked-out the *MIR10B* gene locus in RDEB-SCC1 cells (RDEB-SCC1^*MIR10B*-/-^) using the CRISPR-Cas9 technology, and performed minimal dilution experiments to potentially obtain single clones (Supplementary Fig. S5J,K). In spheroid formation assays we observed, that miR-10b knock-out reduced the stability of aggregates and resulted in an increased number of single cells and fragmented aggregates (Figure 3A-C). While PCR-mediated confirmation of *MIR10B* knock-out showed only bands corresponding to successful deletion, we found that over time single cells that had escaped knock-out and subsequent clearance by minimal dilution returned to dominance, independent of a potential proliferative advantage (Figure 3D,E). This was observed in several clones and over several cultivation passages, pointing towards a potential survival advantage of cells expressing miR-10b. When subjecting these mixed cultures again to our assays, their behavior in tumor spheroid formation resembled that of their parental cells (Figure 3A-C). Another striking difference between parental and *MIR10B-/-* cells was the reduced capacity of miR-10b knock-out cells to grow out of tumor spheroids upon transfer to culture dishes. Spheroids adhered to dishes, and circularly outgrowing cells became visible after 24 hrs in RDEB-SCC1 derived aggregates, and to a much lower extent in respective miR-10b knock-out cells. Again, this was reversed in mixed culture experiments (Figure 3F). This outgrowth pattern was also observed in two out of three HC-cSCC derived spheroid assays (Figure 3G).

**Figure 3:**
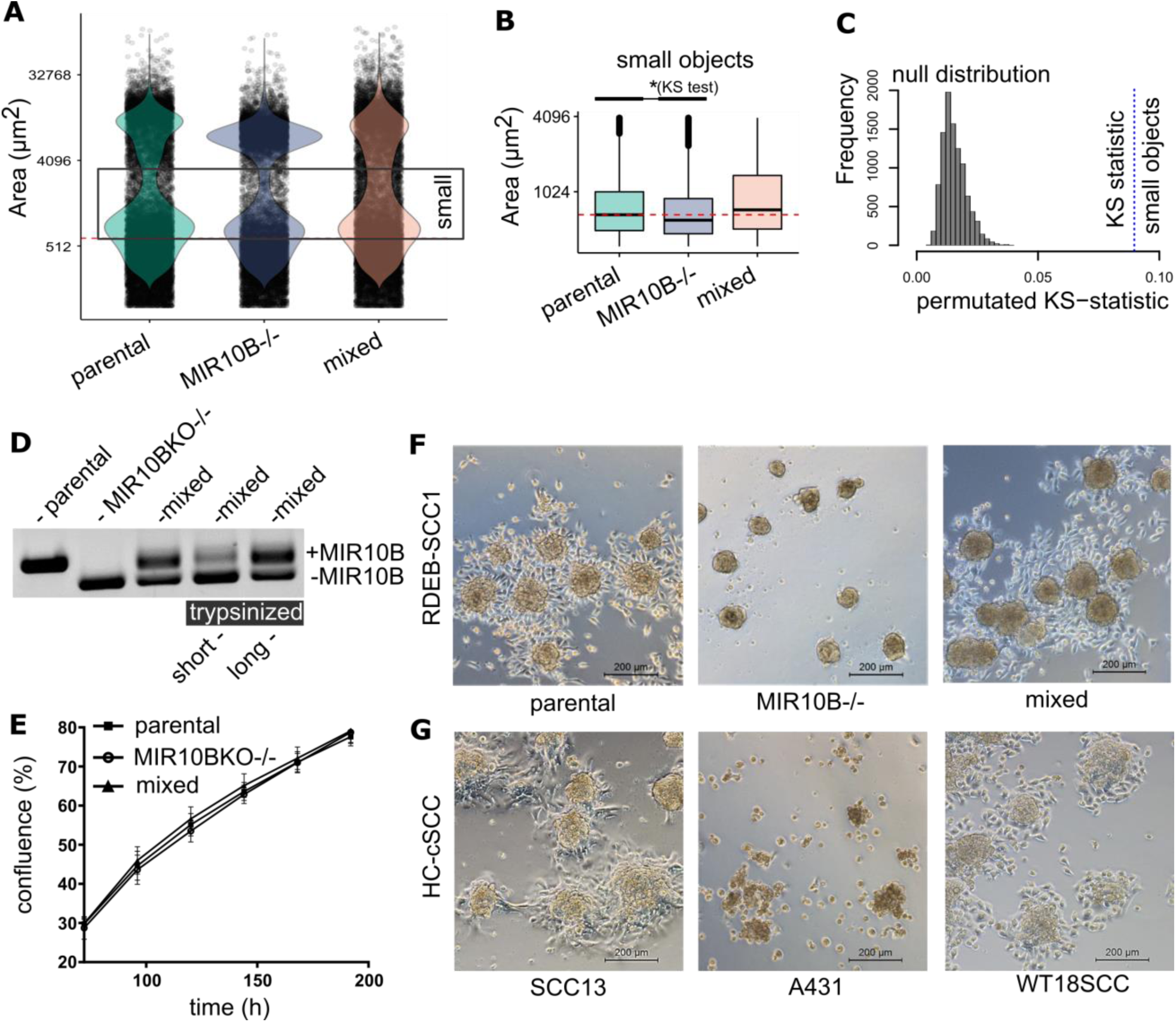
Knock-out of *MIR10B* reduces aggregate sizes. (A,B) Knock-out of *MIR10B* in RDEB-SCC shifts the distribution of cell aggregates towards an increased number of single cells and aggregate fragments (indicated as small objects) in a size distribution analysis by cross-section of formed aggregates compared to parental cells. This effect was reversed in a mixed culture of knock-out and parental cells. (C) The significant difference in size distribution was evaluated by a Kolmogorov-Smirnoff (KS) test, where the null distribution of the KS test statistic was derived by Monte Carlo simulation upon random resampling (parental, MIR10B-/- and mixed). (D) Extended culture period over several passages (>10) of *MIR10B* knock-out cells led to the prevailance of residual miR-10b expressing cells, resulting in a mixed polulation of RDEB-SCC1^MIR10B-/-^ and parental miR-10b expressing cells. In addition, upon short trypsinization of a mixed population, we found that *MIR10B* knock-out cells detached earlier, whereas the remaining cells, showed an enrichment of those cells still harboring the genomic MIR-10b locus. (PCR on *MIR10B* locus in isolated genomic DNA). (E) Proliferation of miR-10b expressing parental cells showed no difference to *MIR10B* knock-out cells or to a mixed population. Proliferation was assessed over 7 days. Mean ± SEM are given, n=4 ind. repeats. (F) Spheroids formed by miR-10b expressing parental RDEB-SCC cells and a mixed population showed outgrowth of cells 24 hrs post aggregate transfer to cell culture plates. This phenomenon was much less pronounced in *MIR10B* knock-out cells. (G) Two out of three HC-cSCC showed outgrowth from spheroids within 24 hrs post transfer similar to RDEB-SCC.

As spheroid formation is a key attribute of cancer stem cells (CSCs), and points towards the presence of cancer stem cell-like properties, we next analyzed accepted CSC markers (CD44 / CD24) (33). RDEB-SCC cultures showed an overall high expression of CD44, and varying levels of CD24, including a CD44^high^ / CD24^-/low^ cell population. In support of the hypothesis that miR-10b expression was associated with a CSC-like phenotype, overexpression of miR-10b in KCs resulted in an overall ∼2-fold enrichment of CD44^high^ / CD24^-/low^ cells (Supplementary Fig. S6B,C).

We next examined whether the above observed properties were attributed to mobility-, or adhesion-associated mechanisms (32). Therefore, we conducted wound closure assays using miR-10b overexpressing (KC^miR-10b^) and *MIR10B* knockout (RDEB-SCC1^*MIR10B-/-*^) cells. We found that those cells expressing high levels of miR-10b (i.e. RDEB-KC^miR-10b^ versus parental, SCC1^MIR10B-/-^ versus parental) demonstrated significantly impaired mobilization (time point 6 hrs: p < 0.01) (Figure 4A,B). In addition, when transiently transfecting SCC1^MIR10B-/-^ with a miR-10b mimic, as well as when using the mixed population, gap closure was accelerated again (Figure 4B,C). No impact on proliferation was observed in the presence or absence of miR-10b.

**Figure 4:**
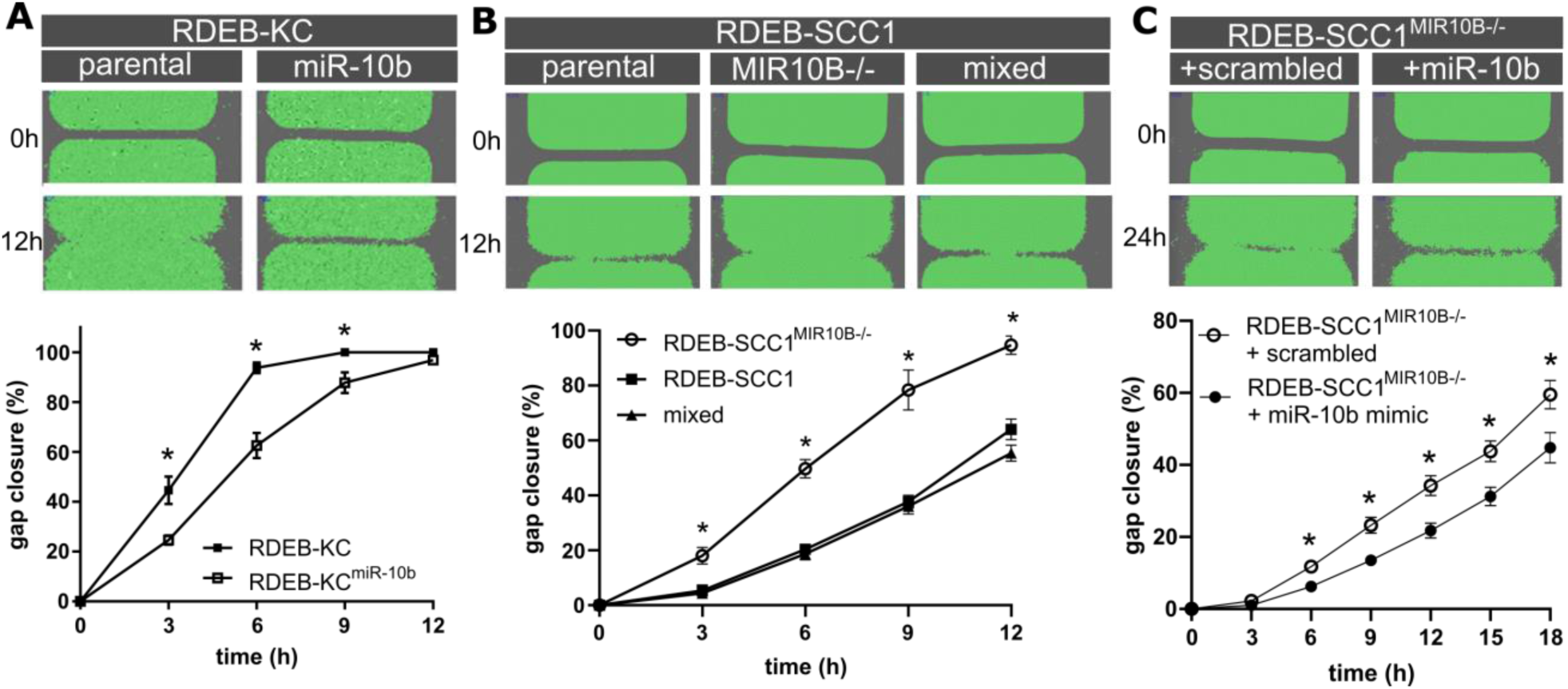
Gap closure is affected by miR-10b expression. (A) Stable overexpression of miR-10b in an RDEB keratinocyte line prolonged the time to gap closure compared to parental cells (RDEB-KC), n≥8 ind. experiments. (B) RDEB-SCC1^MIR10B-/-^ cells showed an enhanced gap closure compared to parental cells (RDEB-SCC1), and a mixed culture (n≥3 ind. Experiments). (C) Transient transfection of miR-10b mimic into RDEB-SCC1^MIR10B-/-^ accelerated gap closure compared to scrambled control (n=2 with 4 technical replicates each). Mean ± SEM is given, * p < 0.05, unpaired t-test.

Taken together, our data suggest for the first time that CSC-like properties can be conferred by miR-10b, a heretofore unknown aspect of miR-10b driven malignancy in cSCCs.

### DIAPH2 is deregulated in RDEB-SCC and predicted to be a target of miR-10b

To analyze the impact of miR-10b on its targetome, we first screened the scientific literature in an automated text-mining approach for miR-10b in cancer, which highlighted the previously reported, direct miR-10b target transcription factor *HOXD10* (14, Supplementary Fig. S7A). We observed significantly reduced levels of HOXD10 protein in three RDEB-SCC cell lines compared to HC-KC, and also in an RDEB-SCC tissue section, compared to HC- and RDEB-skin (34, Supplementary Fig. S7B-D).

To next identify novel downstream targets of miR-10b by the example of RDEB, data driven miR-10b target identification was implemented based on transcriptome data generated from miRNA microarray-matched RDEB-SCC and RDEB-KC samples. Of the 576 differentially expressed genes identified (≥ 2-fold ↓↑) in RDEB-SCCs, 114 were reported in merged repository data (n = 3,923) of validated miR-10 targets (miRTarbase v6.1), as well as computationally predicted targets by seed sequence and evolutionary conservation (TargetScan v7.2). Dysregulated putative miR-10 interaction partners were further prioritized by strength of their inverse correlation of expression with miR-10 signal (Supplementary Fig. S7E). To nominate a disease relevant miR-10b target, we analyzed publicly available survival data from metastatic stage IV head and neck squamous cell carcinoma (HNSCC) (n = 86 patients, The Cancer Genome Atlas / TCGA), as this cancer type was previously described to have high genetic similarities to RDEB-SCC (8). The top 20 candidate miR-10b targets with highest inverse correlation were then used to stratify HNSCC patients (Supplementary Fig. S8A and Supplementary Table S6). We nominated diaphanous related formin 2 (*DIAPH2)* for further evaluation based on significant differences in Kaplan Meier survival curves (p < 0.05, log-rank test, Figure 5A), together with the fact that it was listed as a putative target of miR-10b. Its potential disease-relevance was substantiated by its recent association with colon carcinoma, and its potential role in actin-organization and microtubule stabilization (35-37). When re-analyzing normalized RNA-seq data generated by Cho *et al.*, retrieved from GEO-repository (GSE111582), both, *DIAPH2* and *HOXD10*, showed lower expression in RDEB-SCC versus RDEB-skin tissue (8, Supplementary Fig. S8B,C). *DIAPH2* was also slightly downregulated in KC^miR-10b^ as shown by sqRT-PCR (Supplementary Fig. S5B). However, when analyzing DIAPH2 protein expression in KC^miR-10b^ and RDEB-SCC^MIR10B-/-^ by Western blot analysis and immunofluorescence, results were not conclusive, pointing towards the impact of further regulatory mechanisms influencing DIAPH2 expression (data not shown).

**Figure 5:**
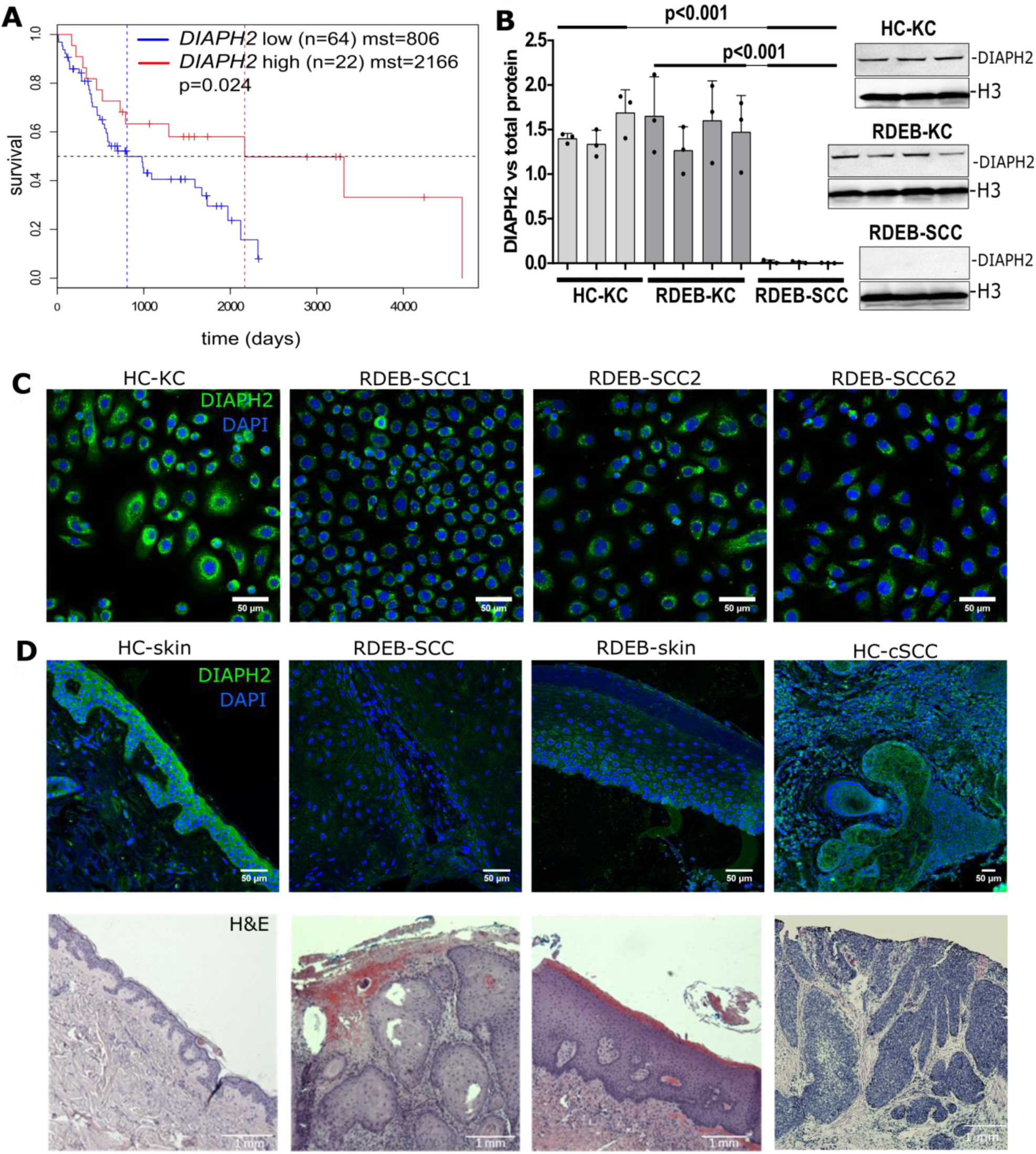
*DIAPH2* is a direct downstream target of miR-10b in RDEB-SCC. (A) Kaplan-Meier curves of the metastatic HNSCC (TCGA RNAseq and clinical data filtered for stage IV, N3). Data show significant (p = 0.024, log-rank test) impact of *DIAPH2* expression (75% quantile is considered for stratification between high and mid/low expression) on mean survival time (mst) mst _DIAPH2_^high^ = 2,166 vs. mst _DIAPH2_^low^ = 806 days. Overall expression of DIAPH2 in cell cultures from experimental groups showed significant downregulation of *DIAPH2* in Western blot analysis (p < 0.001; n = 3; t-test; error bars represent SEM), and immunofluorescence microscopy in cell cutures and (D) tissues.

To confirm *DIAPH2* as a direct target of miR-10b, a dual luciferase reporter assay was established by cloning the 3’UTRs of *DIAPH2* and *HOXD10* as a control, respectively, downstream of a firefly luciferase reporter gene. Constructs were then co-transfected with a miR-10b mimic into HC-KCs, which express only low levels of endogenous miR-10b. In the presence of miR-10b mimic, luciferase signal was significantly reduced (p-value^DIAPH2^ = 0.015; p-value^HOXD10^ < 0.01) compared to scrambled (SCR) control (Supplementary Fig. S7F,G).

We next assessed *DIAPH2* expression in cultured RDEB- and control KC. Both, at the mRNA and the protein level, DIAPH2 was downregulated in RDEB-SCC cells (Supplementary Fig. S7I, Figure 5B,C). Notably, following transient transfection of HC-KCs with miR-10b mimic, DIAPH2 protein levels were found to be significantly reduced (∼29%, p = 0.005) compared to SCR control (Supplementary Fig. S7H). DIAPH2 downregulation was also evident in RDEB-SCC tissue sections (Figure 5D).

To examine if a loss of DIAPH2 in keratinocytes phenocopies the observed migratory behavior in KC^miR-10^, we knocked out *DIAPH2* using the CRISPR/Cas9 technology (1090KC^*DIAPH2-/-*^). Clones were generated by minimal dilution and knock-out was confirmed by Sanger sequencing and Western blot analysis (Supplementary Fig. S5C,D). While proliferation was not altered, *DIAPH2-/-* cells showed significantly impaired motility over HC-KCs (Supplementary Fig. S5E,F). In addition, when analyzing HC-KC^*DIAPH2-/-*^ in a 3D-sphere formation assay, a trend towards enhanced aggregation was observed, with a distribution of spheroid size similar to cSCCs (Supplementary Fig. S5G,H). The functional impact of *DIAPH2* knock-out resembled our observations in miR-10b overexpressing KCs.

Our work taken as a whole has demonstrated that miR-10b is overexpressed in cSCC cells and tissues, and that miR-10b confers tumor-associated properties to KCs. In addition, deregulation of DIAPH2, a novel putative miR-10b downstream target, might represent a contributing factor in cSCC malignancy.

## Discussion

Various factors are presumed to contribute to the development of SCCs in RDEB patients, however, the role of miRNAs in the context of RDEB has not yet been addressed. In this study, our aim was to identify miRNAs involved in development or progression of particularly aggressive cSCCs, by profiling the miRNome of human cSCCs, as well as keratinocytes. As miR-10b was found to be significantly upregulated in cSCC *in vitro* and *in situ* we next examined its functional impact on normal human keratinocytes. Contrary to initial expectations keratinocytes overexpressing miR-10b migrated more slowly and showed no differences in proliferation and / or invasive potential compared to controls. However, the overexpression of miR-10b improved the ability of cells to form stable 3D-spheroids: a capacity for anchorage-independent survival, generally attributed to cells with the potential to initiate tumor growth, such as CSCs-which resembled those derived from aggressive cSCC cells. This shift towards stemness upon miR-10b overexpression in keratinocytes was confirmed through an enrichment of CD44^high^ / CD24^-/low^ cells (38). Unfortunately, tumor initiation experiments in immunodeficient mice did not result in the engraftment of cells, or the development of a tumor, likely due to their overall non-malignant background of RDEB keratinocytes. Notably, not all RDEB-SCC lines are able to successfully engraft in immunodeficient mice, highlighting that other factors, such as a supportive microenvironment, are required.

However, the observation that miR-10b is overexpressed and may have a functional role in mediating stemness does highlight a novel aspect of miR-10b function in cSCC, that is separate from its well demonstrated role in EMT.

While it has been repeatedly reported that EMT is necessary for cells to disseminate from the primary tumor and become circulating tumor cells (CTCs), complementary models of metastasis exist (31, 39). One pre-requisite for CTCs is to maintain their ability to revert from the mesenchymal to the epithelial phenotype (MET). Another is based on the concept of “collective” or “cohort” migration. In this case it is assumed that epithelial-like and mesenchymal-like cancer cells circulate in clusters and act cooperatively to promote metastatic growth, without the need to undergo complete EMT/MET (40). Ultimately, a key event during metastasis is that cells with a tumor-initiating capacity reach and establish at distant sites to form new colonies, which requires cell adhesion (31). We hypothesize that the increased adhesive potential and sphere formation capacity following overexpression of miR-10b, might not only increase cell survival in the circulation, but also facilitate extravasation at new sites, and explains in part the propensity of aggressive cSCCs to metastasize (41).

Alternatively, changes in the actin- and / or microtubule cytoskeleton, which impact cell motility (42), could account for slower wound closure rates and increased stickiness in RDEB skin. In this context, the putative miR-10b target DIAPH2, a protein belonging to the formin homology family, which has previously been associated with actin filament assembly, microtubule formation and vesicle shuttling, might be a modulator of cell motility in RDEB-SCCs (43,44). In this study, DIAPH2 was predicted to be a target of miR-10b *in silico*, which we additionally confirmed using a luciferase-reporter assays. Notably, while miR-10b overexpression did not result in efficient knock-down of DIAPH2 in RDEB-KCs, a significant downregulation of DIAPH2 was observed in RDEB-SCC expressing high endogenous levels of miR-10b. This indicates potential alternative pathways of DIAPH2 regulation in non-malignangt cells. Nonetheless, CRISPR-mediated knock-out of *DIAPH2* resulted in functional outcomes similar to those observed in miR-10b overexpressing keratinocytes, including attenuated migration and an enhanced capacity to form 3D-spheroids. While only limited information on DIAPH2 is available, DIAPH3, a closely related family member with 57 % sequence homology and several common protein binding domains, does promote cell growth and metastasis in hepatocellular carcinoma (45). Supporting our findings, knock-down of DIAPH3 results in reduced migration (46). DIAPH2 is predicted to interact with small RHO-GTPases, which are implicated in motility, and frequently downregulated in different types of SCCs (47,48). In line with this, our microarray data also indicate a slight downregulation of *RHOC* and *RHOD* in RDEB-SCC. Considering this, a similar function of DIAPH2 in the context of SCCs is likely but needs to be further explored. However, while we could show that *DIAPH2* is a potential target of miR-10b by luciferase assays, and expression levels can be correlated to miR-10b expression in patient samples, it seems that also other regulatory networks impact *DIAPH2* expression, as downregulation was not conclusive in miR-10b overexpressing cells. Therefore, regulatory networks affecting DIAPH2 expression and its biological role will need further investigation in the future.

In summary, overexpression of miR-10b impacts cellular processes at various levels. The extent of the response is dependent on the cellular context, the presence of certain target mRNAs, and/or mutations in those target mRNAs (49). In the context of cSCCs, we showed that miR-10b confers anchorage-independent spheroid formation capacities, indicating a phenotypic shift towards stem cell-like properties. We hypothesize, that these findings might be involved in tumor progression at the stages of extravasation and metastatic colonization, processes in which both anchorage-free survival, as well as cell surface interaction, are important pre-requisites.

## Conclusion

Overall, our results demonstrate for the first time the upregulation of miR-10b in aggressive RDEB-SCCs. While new, these findings provide a potential so far unreported pro-metastatic function of miR-10b, as this miRNA confers cancer stem cell-like properties, like increased adhesion, spheroid formation capacities and associated cellular outgrowth, which are essential characteristics associated with metastasis. The results of this study provide an exciting new opportunity for the development of future therapies to treat this rare and debilitating disease. In addition, miR-10b expression in RDEB-SCC might hold potential as a biomarker for early diagnosis and / or therapy monitoring in the context of aggressive cSCC, as these tumors often escape early detection due to their predominant emergence within chronic wounds.

## Notes

Monika Wimmer and Roland Zauner contributed equally to this work.

## List of abbreviations

cSCC: cutaneous squamous cell carcinomas
CSCs: cancer stem cells
CTCs: circulating tumor cells
DIAPH2: Diaphanous Related Formin 2
ECM: extracellular matrix
EMT: epithelial to mesenchymal transition
FDR: false discovery rate
FFPE: formalin-fixed paraffin embedded
GAPDH: Glycerinaldehyd-3-phosphat-dehydrogenase
H&E: Hematoxilin & eosin
HC: healthy control
HNSCC: head and neck squamous cell carcinoma
HOXD: homeobox D
IHC: immunohistochemistry
KC: keratinocytes
LNA: locked-nucleid acid
MET: mesenchymal to epithelial transition
miRNAs: micro-RNAs
onco-miRs: oncogenic miRNAs
PCA: principal component analysis
RDEB: recessive-dystrophic epidermolysis bullosa
LM: lymphnode metastasis
RNA-seq: RNA sequencing
SCC: squamous cell carcinomas
SCR: scrambled
SEM: standard error of mean
SSC: saline-sodium citrate
TCGA: The Cancer Genome Atlas
TUBA1: tubulin alpha 1

## Declarations

### Ethics approval and consent to participate

RDEB tissue samples collected in Salzburg were obtained from patients undergoing dermatological surgery upon written, informed consent. Ethical approval was granted by the ethics committee of the county of Salzburg (vote number: 415-EP/73/192-2013) for RDEB and 1090KC cell isolations. For the generation of “SKC”-HC cell lines human full-thickness skin was obtained from biologic waste material during plastic reconstructive surgery after informed consent as approved by the ethical committee of the county of Salzburg (vote number: 415-E/1990/8–216). Skin was processed as described previously (50).

### Consent for publication

Not applicable.

### Availability of data and material

The data that support the findings of this study were submitted to the Gene Expression Omnibus Database (Accession: GSE130767 for miRNA and GSE130925 for transcriptome microarray data). And the data are available from https://www.ncbi.nlm.nih.gov/geo/query/acc.cgi?acc=GSE130767 and https://www.ncbi.nlm.nih.gov/geo/query/acc.cgi?acc=GSE130925

### Authors’ contributions

MW and RZ performed experiments, designed and generated data processing workflows, interpreted the experimental results and assisted in conception and writing of the manuscript. MA participated in the qRT-PCR, immunostaining, western blots, FACS and assisted in data interpretation. JPH participated in FACS analysis and assisted in manuscript writing. CGG, MR, SH, EMM, DS, TK participated in isolating primary cells, cloning of retroviral vectors and generation of cell lines. NN and JP performed miRNA sequencing. MS, PDS and ASM assisted in the optimization of the miRNA ISH protocol. TL, JWB, JR and ASM contributed in writing and reviewing the manuscript. VW conceived and designed this study, interpreted the data and wrote the manuscript. All authors reviewed the manuscript and approved the submission.

## Acknowledgements

We would like to thank Prof. Andrew South and Prof. Leena Bruckner-Tuderman for providing cell lines RDEB-SCC1 and RDEB-SCC2, and Tzipi Cohen Hyams, PhD and A/Prof Murray Killingsworth, Ingham Institute for Applied Medical Research, Sydney for their technical support. In addition we would like to thank the group of Prof. Herbert Reitsamer, University Clinic of Ophthalmology and Optometry / Paracelsus Medical University Salzburg, for providing access to their confocal laser-scanning unit (Axio Observer Z1 attached to LSM710, Zeiss). Finally, Andrea Wagner from the Institute of Tendon & Bone Regeneration, Spinal Cord Injury and Tissue Regeneration Center Salzburg (SCI-TReCS), Paracelsus Medical University, for her support in IHC-tissue processing.

## Additional Files

*File name:*

AdditionalFile1.pdf

*Description of Data:*

Additional information on cell lines, bioinformatic data processing, CRISPR/Cas9-mediated knock-out of DIAPH2, Flow cytometry for CD44 and CD24 expression, miRNA library preparation and next generation sequencing, Luciferase-reporter assay.

Supplementary data summarized in tables: cell lines, antibodies, primers sequences, top 30 miRNAs upregulated in RDEB-SCC versus RDEB-KCs, miR-10b target selection and validation.

Supplementary experimental data in figures: generation of experimental cell lines, miR-10b target identification and selection, miRNA probe specificity for in situ hybridization on cultured cells and single cell resolution image-based data analysis workflow, miRNA probe specificity for in situ hybridization on cultured cells, controls for miRNA probe specificity in situ hybridization on FFPE tissue sections, characterization of experimental cell lines, spheroid formation assay, miR-10b target selection and validation, survival analysis and target evaluation.

