## Supplemental Information for "MiRNA-10b marks aggressive squamous cell carcinomas, and confers a cancer stem cell-like phenotype"

Monika Wimmer[1]*, Roland Zauner[1]*, Michael Ablinger[1], Josefina Piñón-Hofbauer[1], Christina Guttmann-Gruber[1], Manuela Reisenberger[1], Thomas Lettner[1], Norbert Niklas[2], Johannes Proell[3], Mila Sajinovic[4], Paul De Souza[4,5], Stefan Hainzl[1], Thomas Kocher[1], Eva M. Murauer[1], Johann W. Bauer[1,6], Dirk Strunk[7], Julia Reichelt[1], Albert S. Mellick[4,8], Verena Wally[1#]

[1]Research Program for Molecular Therapy of Genodermatoses, EB House Austria, Department of Dermatology and Allergology, University Hospital of the Paracelsus Medical University, 5020 Salzburg, Austria.

[2] Red Cross Transfusion Service of Upper Austria, 4020 Linz, Austria.

[3] Center for Medical Research, Medical Faculty, Johannes-Kepler-University, 4020 Linz, Austria.

[4] Medical Oncology, Ingham Institute for Applied Medical Research, Liverpool NSW 2170, Australia.

[5] School of Medicine, Western Sydney University, Campbelltown, NSW 2560, Australia.

[6]Department of Dermatology and EB House Austria, University Hospital of the Paracelsus Medical University Salzburg, 5020 Salzburg, Austria.

[7]Institute of Experimental & Clinical Cell Therapy, Paracelsus Medical University, 5020 Salzburg, Austria.

[8] School of Medicine, University of New South Wales, Kensington NSW 2052, Australia.

*MW and RZ equally contributed to the work.*

### Supplementary online content

**Supplementary Information**

Cell lines

Bioinformatic data processing

CRISPR/Cas9-mediated knock-out of *DIAPH2*

Flow cytometry for CD44 and CD24 expression

miRNA library preparation and next generation sequencing

Luciferase-reporter assay

**Supplementary Tables**

Supplementary Table 1: Cell lines.

Supplementary Table 2: Antibodies.

Supplementary Table 3: Primers sequences.

Supplementary Table 4: Top up- and downregulated miRNAs in microarray analysis: RDEB-SCC versus RDEB-KCs.

Supplementary Table 5: Top up- and downregulated miRNAs in microarray analysis: RDEB-SCC versus RDEB-KCs.

MiR-10b target selection and validation.

**Supplementary Figures**

Supplementary Figure S1: MiR-10b target selection and validation.

Supplementary Figure S2: Controls for miRNA probe specificity in *in situ* hybridization on FFPE tissue sections.

Supplementary Figure S3: MiRNA probe specificity for *in situ* hybridization on cultured cells and single cell resolution image-based data analysis workflow.

Supplementary Figure S4: MiRNA probe specificity for *in situ* hybridization on cultured cells.

Supplementary Figure S5: Generation of experimental cell lines.

Supplementary Figure S6: Characterization of experimental cell lines.

Supplementary Figure S7: Validation of miR-10b targets HOXD10 and DIAPH2.

Supplementary Figure S8: Survival analysis and target evaluation.

#### Supplementary references

#### Supplementary Information

##### Cell lines

In this study we utilized cancer cells derived from primary skin tumors of RDEB, otherwise healthy, non-RDEB patients, as well as RDEB- and healthy control keratinocytes (HC-KCs). All RDEB cancer patients have in the meantime succumbed to the metastatic disease, underlining the devastating consequences of this genetic condition. Cell lines used for non-RDEB cSCCs have diverse characteristics. A431 (ATCC, CRL-1555) were derived from an epidermoid carcinoma from a 85 year old female patient and express high levels of epidermal growth factor receptor (EGFR) (1). When injecting A431 cells into mice, in one out of six mice a metastatic lesion was detected (2).

SCC13 were generated by Rheinwald *et al*. from a facial lesion of a female 56-year old female patient, who had received radiotherapy before excision (3). While we have no knowledge about the fate of the patient, SCC13 cells have been shown to be able to metastasize in a mouse xenograft model (4).

WT18SCC were isolated from a facial primary, moderately differentiated SCC of a female 66-year old patient. In the course of histology it was found that a BCC was associated at a tangential section plane. Because of that, it can not be excluded, that a minor proportion of BCC cells are present in the cell cultures. In a second surgery, histology also revealed presence of an actinic keratosis, which could not be removed in healthy complete. However, during the patient's follow-up, lymph node metastasis occurred which were treated by complete dissection. The patient has now been tumor-free for about 3 years (Supplementary Table 1).

##### Bioinformatic data processing

MiRNome analysis and microarray raw data (.CEL files) were pre-processed using the robust multi-array average (RMA, oligo package, v1.46.0) background correction, normalization, log2-transformation and annotated with miRNA-4_1-st-v1 configuration file. Unsupervised clustering was performed using principal component analysis (PCA, FactoMineR v1.41) and factor loadings were used to identify miRNAs as drivers of experimental group separation in dimension 1. Differential expression of batch effect corrected (SVA package v3.30.0) miRNA with expression log2(signal) > 2.5 in at least 3 samples was assessed via Linear Models for Microarray Data (limma, v3.38.3). Overrepresentation analysis (clusterProfiler, v3.10.1) of miR-10a/b targetome (miRTarbase v.6.1) was performed on MSigDB hallmark gene sets (v6.1 Broad Institute, Massachusetts Institute of Technology) and significance ($p \leq$

0.05; hypergeometric test) of enriched genesets were presented in a radial plot for either miR-10a, miR-10b or shared targets.

Data driven miR-10a/b target identification – transcriptome microarray raw data (.CEL files) were pre-processed by RMA (oligo package, v1.46.0) background correction, normalization, log2-transformation and annotated with Clariom_D_Human.na36.hg38.transcript file. Greater than 2-fold de-regulated mRNAs (limma) in RDEB-SCC versus RDEB-KC were matched to union of miR-10a and miR-10b targets reported in TargetScan (v7.2) and miRTarbase (v6.1). The resulting predicted target list was prioritized by strength of inverse correlation (Pearson correlation) between mean of miR-10a/b (correlation of microarray log2 miRNA expression signals with TaqMan qPCR $\Delta$Cq values of respective miR-10 family members supports the preferential use of average miR-10 microarray signals, Supplementary Fig. S7E) and target mRNA log2-transformed, normalized expression signals. Final candidate selection was based on investigating the impact of miR-target gene expression on survival time of head and neck SCC (HNSCC) patients. HNSCC RNAseq and clinical data was retrieved from The Cancer Genome Atlas (TCGA, RTCGAToolbox, v2.12.1) and filtered for tumor stage IV and reports of lymphnode metastasis (N3 - TNM tumor staging). Patients were stratified by expression levels of prior short-listed miR-target genes into high and mid/low expression groups (75% quantile cut-off) and difference in survival time curves was tested applying log-rank test (survival, 2.43-3).

For scientific literature driven miR-10b target identification, the search term “(cancer) AND (miR-10b)” was used to screen (RISmed, v2.1.7) PubMed articles. Biological terms were extracted from abstract text bodies through named entity recognition by accessing geniatagger (v3.0, School of Computer Science, University of Manchester). Terms were summarized and highlighted by frequency of their occurance in abstracts (wordcloud, v0.2.1).

For *DIAPH2* expression analysis in RDEB-SCC and RDEB-skin tissue, normalized RNAseq data generated by Prof. Andy South’s lab (Thomas Jefferson University, USA) was retrieved from GEO repository (GEOquery, v2.50.5, GSE111582) and filtered for *DIAPH1,2,3* and *HOXD10* log2-transformed expression signals.

Data evaluation and analysis of miRNA-seq data was performed via R package DESeq2. Therefore read counts were loaded from .csv files into the statistical program language R (v 3.3.1) for all further processing and data analysis steps. Multiple annotations deriving from 2 bp mismatches were resolved for final read count summary. Annotation codes were adapted for further database screenings, like adding the prefix "hsa-". Normalization and inference testing with fold change estimation was performed via R package "DEseq2" (v 1.12.3). Results were reported in box and dotplots created by "ggplot2" (v 2.1.0).

**CRISPR/Cas9-mediated knock-out of DIAPH2**

DIAPH2 knock-out cell lines were generated using the U6gRNA-Cas9-2A-RFP vector (Sigma Aldrich, HS000336959), including a guide RNA specifically targeting *DIAPH2* exon 1 (Supplementary Table S3). Upon Xfect-mediated transfection of 1090HC-KC, efficiency in the bulk was analyzed using a T7-endonuclease I (New England Biolabs, E3321), according to the manufacturer's instruction and as describe elsewhere (5). Primers were designed to flank the target region, and which will produce two fragments of different sizes upon T7-endonuclease I digest (Supplementary Table S3). Clones were generated by minimal dilution and knock-out was confirmed by Sanger sequencing and Western blot analysis and semi-quantitative (sq)RT-PCR, as described.

**Flow cytometry for CD44 and CD24 expression.**

$10^6$ cells were re-suspended in PBS and double stained with CD44-FITC (FITC anti-mouse/human CD44, clone:IM7, Cat.103005, Lot.B228503) and CD24-PE (PE anti human CD24, clone:ML5, BioLegend, Cat.311105, Lot.B216880). Cytometry was performed on a Gallios Flow Cytometer (Beckman Coulter) and data were analysed using the Kaluza analysis software (Beckman Coulter). Isotype controls were included to exclude non-specific staining (PE Mouse IgG2a, k isotype ctr., clone:MOPC-173, Cat.400211, Lot.B265821; FITC Rat IgG2b, k isotype ctr., clone:RTK4530, Cat.400605, Lot.B265825).

**miRNA library preparation and next generation sequencing**

*miRNA library preparation*

cDNA libraries from ~50 ng AGO2 co-immunoprecipitated small RNA pools were generated using the NEBNext Multiplex Small RNA Library Prep Set for Illumina (Set2; NEB #E7580S/L) according to the manufacturer's protocol, including quality control and polyacrylamide gel-based size selection steps. The cDNA libraries were purified using the "QIAquick PCR purification kit" (Qiagen, Cat No./ID: 2810). Concentrations of miRNA and cDNA were analyzed using a Qubit fluorometer and quality controls of cDNA libraries were done using an Experion™ Automated electrophoresis system (Bio-Rad) using the "Experion DNA 1K analysis kit" (Experion, #700-7107) following the manufacturer' instruction manual.

*miRNA next generation sequencing*

All libraries were subtracted to Illumina next-generation sequencing (NGS) on an Illumina MiSeq platform. For the generation of the final sequencing libraries, up to four samples were simultaneously equimolarly pooled. Molarities for pooling were calculated by fragment size (Agilent) and quantification using PicoGreen (Thermo Fisher, P11496). The final concentration of the sequencing libraries were 12 pM in which 1% depicted the spike-in of PhiX sequencing control. Sequencing itself was performed according to the manufacturer' instruction by using a 50-cycle v2 MiSeq kit (Illumina, MS-102-3001). The sequencing pipeline included base calling and demultiplexing, generating a total of 92.2 M reads (in six runs) with Q30 ranging from 90.6 % to 97.1 % and an average error rate of 0.33 % (measured by PhiX control). Adapters contained in the sample sequences were trimmed after sequencing, using the CLC Genomics Workbench 7 (Qiagen). After trimming, the size distribution showed a typical miRNA pattern (around 22-25 bp), no additional filters were applied. The trimmed sequences were counted and grouped (by equal bp). Annotated sequences were identified by miRBase 21 and reported according to the database annotation. Mature length variants were determined by using the default settings of the "Annotate and Merge Counts" (2 bp tolerance). Reports included miRNA name and maturity as well as their abundance. Unmatched sequences (without annotation) were mapped against the whole human genome (using default parameters). Results were provided in excel and comma-separated-value (.csv) files, including all sequences of identified RNAs and the respective annotation if available.

**Luciferase-reporter assay**

For Dual-Luciferase® Reporter (DLR™) assay system (Promega, E1910) 3'UTRs of *DIAPH2* and *HOXD10*, respectively, were cloned into the pmirGLO Dual-Luciferase miRNA Target Expression Vector (Promega), downstream the firefly luciferase coding region. For primers see Supplementary Table S3. 5 µg of reporter vector were co-transfected with 100 pmol miR-10b-5p mimic, or a scrambled control (SCR) into primary HC-KCs at ~70 % confluency in a 6-well format. Cells were lysed in passive lysis buffer after 24 hrs and luminescence was measured using the Tecan Spark 10M plate reader. Luciferase signals were normalized to Renilla signals for comparison of co-transfection with miR-10b or SCR.

#### Supplementary Tables

##### Supplementary Table S1: Cell lines.

| | age (yrs) | sex | *COL7A1* mutation (allele 1 / allele 2) | localization | tumor type / history of metastasis / metastatic potential | reference |
|---|---|---|---|---|---|---|
| **1090KC** | 23 | m | healthy control | arm | - | - |
| **SKC013** | 35 | f | healthy control | abdominal | - | - |
| **SKC015** | 36 | f | healthy control | abdominal | - | - |
| **SKC018** | 31 | m | healthy control | abdominal | - | - |
| **1102KC** | 38 | f | healthy control | lower leg | - | (6), (NHK 3) |
| **RDEB-01KC** | 20 | f | c.2005C>T; p.R669X | leg | - | - |
| **RDEB-03KC** | 18 | m | c.425A>G / c.5440G>T p.K142R / p.R1814C | right inner upper arm | - | - |
| **RDEB-29KC** | 2 | m | c.425A>G / c.520G>A p.L142R / p.G174R | foreskin | - | - |
| **RDEB-30KC** | 2 | m | c.425A>G / c.520G>A p.L142R / p.G174R | foreskin | - | - |
| **RDEB-43KC** | 17 | f | c.4027C>T / c.425A>G p.R1343X / p.K142R | right inner thigh | - | - |
| **RDEB-53KC** | 32 | f | c.2005C>T, homozygous | unknown | - | - |
| **RDEB-55KC** | 21 | m | c.976+4A>C p.INS34AA | left upper thigh | - | - |
| **RDEB-57KC** | 6 | w | c.427-2A>G / c.4172insC; p. splice site / p.1391fsX10 | unknown | - | - |
| **RDEB-SCC1** | 32 | f | c.8244dupC homozygous | SCC shoulder | unknown | (7), (RDEB6) |
| **RDEB-SCC2** | 54 | m | c.3832-1 G>A / undetermined | SCC hand | primary, poorly differentiated, development of metastatic disease | (7), (RDEB3) |
| **RDEB-SCC53** | 31 | f | c.2005C>T homozygous | SCC elbow | primary, development of metastatic disease | - |
| **RDEB-SCC62** | 29 | f | c.682+1G>A / c.7474C>T p.R2492X | SCC lower arm | primary, highly differentiated, development of metastatic disease | (5) |
| **A431** | 85 | f | otherwise healthy | cSCC | primary, metastasize in mouse model | ATCC |
| **SCC13** | 56 | f | otherwise healthy | cSCC facial | radiation before removal, metastasize in mouse model | (3) |
| **WT18SCC** | 66 | f | otherwise healthy | cSCC facial with tangential BCC | primary, moderatly differentiated, lymph node metastasis | - |
| **DDEB-22KC** | 46 | m | p.G2043R | right inner upper arm | - | - |
| **DDEB-25KC** | 25 | m | c. 6127G>A/ homozygous p.G2042R | left inner upper arm | - | - |

Several cell lines from different donors were used, including RDEB-SCC (n = 4), HC-cSCC (n = 3), RDEB-KC (n = 8), DDEB-KC (n = 2) and HC-KCs (n = 5) lines. Patients' age, gender, and - where applicable - tumor stadium, history of metastasis of metastatic potential in a mouse model, as well as tumor localization are given. For RDEB-patients, respective *COL7A1* mutations are included.

**Supplementary Table S2: Antibodies.**

| antigen / host species | catalog number / Lot | clone no | company | reference |
|---|---|---|---|---|
| **Antibodies for Western blot analysis** | | | | |
| **primary antibodies** | | | | |
| DIAPH2 / rabbit | HPA005647<br>Lot A37866 | polyclonal | Sigma Life Sciences | (8) |
| TWIST1 / rabbit | T6451<br>Lot 035M4793V | polyclonal | Sigma Life Sciences | (15) |
| Histone 3 / rabbit | PA5-16183<br>Lot SB2344305B | polyclonal | Thermo Fisher Scientific | (9) |
| Annexin 1 / mouse | sc-12740<br>Lot J1216 | EH17a | Santa Cruz | (10) |
| **secondary antibodies** | | | | |
| HRP-labelled goat-α-rabbit IgG2b | K4010<br>Lot 10072150 | | Dako Envision | |
| HRP-labelled goat-α-mouse IgG2b | K4004<br>Lot 10076454 | | Dako Envision | |
| **Antibodies for immunofluorescence microscopy** | | | | |
| **primary antibodies** | | | | |
| DIAPH2 / mouse | sc-55539<br>Lot B0717 | B-11 | Santa Cruz | (11) |
| HOXD10 / rabbit | sc-66926<br>Lot A0510 | polyclonal | Santa Cruz | (12) |
| CD31 / rabbit | PA5-16301<br>Lot NJ1608523 | polyclonal | Thermo Fisher Scientific | (13) |
| CD45 / mouse | MAB1430<br>Lot ILP071808A | 2D1 | R&D Systems | (14) |
| TWIST1 / rabbit | T6451<br>Lot 035M4793V | polyclonal | Sigma Life Sciences | (15) |
| Pan-KRT 14,15,16,19 (A647) / mouse | 563648<br>Lot 8003894 | KA4 | BD Pharmingen | (16) |
| **secondary antibodies** | | | | |
| Alexa Fluor®488 goat-α-mouse IgG (H+L) | A11001<br>Lot 1939600 | polyclonal | Thermo Fisher Scientific | (17) |
| Alexa Fluor®594 goat-α-rabbit IgG (H+L) | A11037<br>Lot 1608397 | polyclonal | Thermo Fisher Scientific | (18) |
| Alexa Fluor®488 goat-α-rabbit IgG (H+L) | A11008<br>Lot 1829920 | polyclonal | Thermo Fisher Scientific | (19) |

**Supplementary Table S3: Primer sequences.**

| | |
|---|---|
| **Primers for sqRT-PCR** | |
| GAPDH_forward | 5’ GCCAACGTGTCAGTGGTGGA 3’ |
| GAPDH_reverse | 5’ CACCACCCTGTTGCTGTAGCC 3’ |
| TUBA1_forward | 5’ATGGAGCCCTGAATGTTGAC3’ |
| TUBA1_reverse | 5’ CTCAAAGCAAGCATTGGTGA 3’ |
| DIAPH2_forward | 5’ GCAGATGATGTGCGTGACCG 3’ |
| DIAPH2_reverse | 5’ GTTAAGGTTCATGTCCTCCATCATTTTTTCAAAGAG 3’ |
| HOXD10_forward | 5’ CCGCAGCTCTCCGCT 3’ |
| HOXD10_reverse | 5’ TCAGACTTGATTTCCTCTTTGCTTTCCTTC 3’ |
| **Primers for cloning** | |
| HOXD10 3’UTR_forward | 5’ GAGCTCGTCTGAGGCCGGT 3’ |
| HOXD10 3’UTR_reverse | 5’ GTCGACGAACTCATTTCCAGAG 3’ |
| DIAPH2 3’UTR forward | 5’ GAGCTCTTCCTGATGCCAAAG 3’ |
| DIAPH2 3’UTR reverse | 5’ GTCGACTAACTTTTACTCCCTA 3’ |
| miR10b_EcoRI_fw | 5’ GATCGAATTCCCGAGGTTGTAACGTTGTC 3’ |
| miR10b_XhoI_rv | 5’ GATCCTCGAGCAAAAATGAAGTTTTTG 3’ |
| **Primers for T7 endonuclease I assay** | |
| DIAPH2genEx1_fw | 5’ CTAGTTCCCTCCCTGTTCTCGCTG 3’ |
| DIAPH2genEx1_rv | 5’ CGTGTTCAATGGCAGGAGGATCCC 3’ |
| **gRNAs for CRISPR** | |
| gRNA1_MIR10B_fw | 5´-TAATACGACTCACTATAGTGAGGTACCTAGGTCGCTGC-3´ |
| gRNA1_MIR10B_rv | 5´-TTCTAGCTCTAAAACGCAGCGACCTAGGTACCTCA-3´ |
| gRNA2_MIR10B_fw | 5´-TAATACGACTCACTATAGCTAGTCTCCATGTCGCACTT-3´ |
| gRNA2_MIR10B_rv | 5´-TTCTAGCTCTAAAACAAGTGCGACATGGAGACTAG-3´ |
| gRNA_DIAPH2 | 5’-GCAGCGAGGAACCCGGTGG-3’ |

**Supplementary Table S4: Top up- and downregulated miRNAs in microarray data analysis.**

**RDEB-SCC (n=4) versus RDEB-KCs (n=6)**

| miRNA | log2(FC) | <log2(Signal)> | p-value | FDR | miRBase Sequence |
|---|---|---|---|---|---|
| hsa-miR-10a-5p | 3.15 | 2.75 | 3.09e-07 | 1.57e-04 | UACCCUGUAGAUCCGAAUUUGUG |
| hsa-miR-146a-5p | 2.99 | 2.32 | 4.86e-03 | 5.66e-02 | UGAGAACUGAAUUCCAUGGGUU |
| hsa-miR-193a-3p | 2.04 | 2.51 | 1.30e-03 | 2.89e-02 | AACUGGCCUACAAAGUCCCAGU |
| hsa-miR-4485 | 1.99 | 5.23 | 7.75e-03 | 7.07e-02 | UAACGGCCGCGGUACCCUAA |
| hsa-miR-194-5p | 1.74 | 4.04 | 3.20e-04 | 1.03e-02 | UGUAACAGCAACUCCAUGUGGA |
| hsa-miR-941 | 1.71 | 3.04 | 2.52e-04 | 1.03e-02 | CACCCGGCUGUGUGCACAUGUGC |
| hsa-miR-3929 | 1.70 | 2.06 | 1.40e-02 | 1.02e-01 | GAGGCUGAUGUGAGUAGACCACU |
| hsa-let-7d-3p | 1.68 | 3.04 | 2.33e-04 | 1.03e-02 | CUAUACGACCUGCUGCCUUUCU |
| hsa-miR-421 | 1.50 | 4.60 | 2.49e-03 | 4.16e-02 | AUCAACAGACAUUAAUUGGGCGC |
| hsa-miR-34c-5p | 1.48 | 2.92 | 1.73e-02 | 1.18e-01 | AGGCAGUGUAGUUAGCUGAUUGC |
| hsa-miR-1973 | 1.45 | 2.59 | 3.23e-03 | 4.58e-02 | ACCGUGCAAAGGUAGCAUA |
| hsa-miR-324-5p | 1.42 | 5.90 | 1.75e-03 | 3.23e-02 | CGCAUCCCCUAGGGCAUUGGUGU |
| hsa-miR-491-5p | 1.42 | 2.52 | 1.63e-03 | 3.23e-02 | AGUGGGGAACCCUUCCAUGAGG |
| hsa-miR-15a-5p | 1.40 | 4.86 | 7.23e-03 | 6.92e-02 | UAGCAGCACAUAAUGGUUUGUG |
| hsa-miR-99a-5p | 1.39 | 4.07 | 1.70e-02 | 1.18e-01 | AACCCGUAGAUCCGAUCUUGUG |
| hsa-miR-619-5p | 1.38 | 4.15 | 6.98e-03 | 6.84e-02 | GCUGGGAUUACAGGCAUGAGCC |
| hsa-miR-139-5p | 1.36 | 2.29 | 3.38e-02 | 1.69e-01 | UCUACAGUGCACGUGUCUCCAGU |
| hsa-miR-23b-5p | 1.34 | 3.01 | 2.62e-02 | 1.47e-01 | UGGGUUCCUGGCAUGCUGAUUU |
| hsa-miR-34c-3p | 1.31 | 3.37 | 4.28e-02 | 1.93e-01 | AAUCACUAACCACACGGCCAGG |
| hsa-miR-132-3p | 1.26 | 5.74 | 7.01e-03 | 6.84e-02 | UAACAGUCUACAGCCAUGGUCG |
| hsa-miR-212-3p | 1.26 | 1.96 | 3.34e-03 | 4.58e-02 | UAACAGUCUCCAGUCACGGCC |
| hsa-miR-769-5p | 1.23 | 2.28 | 1.35e-03 | 2.89e-02 | UGAGACCUCUGGGUUCUGAGCU |
| hsa-miR-501-5p | 1.18 | 3.03 | 3.46e-03 | 4.64e-02 | AAUCCUUUGUCCCUGGGUGAGA |
| hsa-miR-188-5p | 1.18 | 3.08 | 1.42e-02 | 1.02e-01 | CAUCCCUUGCAUGGUGGAGGG |
| hsa-miR-425-3p | 1.16 | 3.48 | 4.56e-03 | 5.56e-02 | AUCGGGAAUGUCGUGUCCGCCC |
| ***hsa-miR-10b-5p*** | ***1.15*** | ***2.31*** | ***2.24e-02*** | ***1.37e-01*** | ***UACCCUGUAGAACCGAAUUUGUG*** |
| hsa-miR-140-3p | 1.15 | 5.02 | 1.02e-02 | 8.54e-02 | UACCACAGGGUAGAACCACGG |
| hsa-miR-342-5p | 1.12 | 2.39 | 1.01e-02 | 8.54e-02 | AGGGGUGCUAUCUGUGAUUGA |
| hsa-miR-192-5p | 1.12 | 2.80 | 3.47e-04 | 1.03e-02 | CUGACCUAUGAAUUGACAGCC |
| hsa-miR-4749-5p | 1.10 | 4.33 | 2.37e-02 | 1.40e-01 | UGCGGGGACAGGCCAGGGCAUC |
| hsa-miR-6124 | 1.07 | 4.32 | 1.28e-02 | 9.68e-02 | GGGAAAAGGAAGGGGGAGGA |
| hsa-miR-151b | 1.05 | 6.32 | 5.12e-05 | 4.09e-03 | UCGAGGAGCUCACAGUCU |
| hsa-miR-671-3p | 1.04 | 3.35 | 1.33e-02 | 9.87e-02 | UCCGGUUCUCAGGGCUCCACC |
| hsa-miR-877-5p | 1.03 | 3.87 | 4.11e-02 | 1.88e-01 | GUAGAGGAGAUGGCGCAGGG |
| hsa-miR-3613-3p | -1.00 | 7.44 | 1.87e-04 | 1.00e-02 | ACAAAAAAAAAAGCCCAACCCUUC |
| hsa-miR-4532 | -1.05 | 6.12 | 2.67e-02 | 1.47e-01 | CCCCGGGGAGCCCGGCG |
| hsa-miR-4674 | -1.07 | 6.43 | 6.23e-03 | 6.55e-02 | CUGGGCUCGGGACGCGCGGCU |
| hsa-miR-205-5p | -1.10 | 12.07 | 8.65e-03 | 7.60e-02 | UCCUUCAUUCCACCGGAGUCUG |
| hsa-miR-200b-3p | -1.16 | 5.09 | 3.77e-02 | 1.79e-01 | UAAUACUGCCUGGUAAUGAUGA |
| hsa-miR-99b-3p | -1.23 | 4.38 | 2.25e-02 | 1.37e-01 | CAAGCUCGUGUCUGUGGGUCCG |
| hsa-miR-296-3p | -1.27 | 3.26 | 2.37e-02 | 1.40e-01 | GAGGGUUGGGUGGAGGCUCUCC |
| hsa-miR-1298-3p | -1.27 | 2.61 | 6.04e-03 | 6.47e-02 | CAUCUGGGCAACUGACUGAAC |
| hsa-let-7e-5p | -1.30 | 9.30 | 8.09e-03 | 7.22e-02 | UGAGGUAGGAGGUUGUAUAGUU |
| hsa-miR-193b-5p | -1.35 | 5.55 | 7.18e-04 | 1.83e-02 | CGGGGUUUUGAGGGCGAGAUGA |
| hsa-miR-424-3p | -1.36 | 7.80 | 2.77e-03 | 4.24e-02 | CAAAACGUGAGGCGCUGCUAU |
| hsa-miR-200a-5p | -1.54 | 4.00 | 4.13e-03 | 5.28e-02 | CAUCUUACCGGACAGUGCUGGA |
| hsa-miR-125a-5p | -1.61 | 9.14 | 1.11e-02 | 8.79e-02 | UCCCUGAGACCCUUUAACCUGUGA |
| hsa-miR-200b-5p | -1.70 | 5.03 | 1.45e-03 | 2.99e-02 | CAUCUUACUGGGCAGCAUUGGA |
| hsa-miR-99b-5p | -1.94 | 8.49 | 2.33e-02 | 1.40e-01 | CACCCGUAGAACCGACCUUGCG |
| hsa-miR-34a-5p | -2.26 | 5.51 | 6.93e-03 | 6.84e-02 | UGGCAGUGUCUUAGCUGGUUGU |

**Supplementary Table S5: Top up- and downregulated miRNAs in microarray data analysis.**

**HC-cSCC (n=3) versus HC-KCs (n=4)**

| miRNA | log2(FC) | <log2(Signal)> | p-value | FDR | miRBase Sequence |
|---|---|---|---|---|---|
| hsa-miR-139-5p | 2.72 | 2.29 | 6.87e-04 | 2.05e-02 | UCUACAGUGCACGUGUCUCCAGU |
| hsa-miR-4485 | 2.55 | 5.23 | 3.16e-03 | 4.13e-02 | UAACGGCCGCGGUACCCUAA |
| hsa-miR-877-5p | 2.28 | 3.87 | 3.39e-04 | 1.40e-02 | GUAGAGGAGAUGGCGCAGGG |
| hsa-miR-126-3p | 2.06 | 3.31 | 9.37e-03 | 6.79e-02 | UCGUACCGUGAGUAAUAAUGCG |
| hsa-miR-1285-3p | 1.99 | 1.99 | 2.94e-07 | 3.94e-05 | UCUGGGCAACAAAGUGAGACCU |
| hsa-miR-27b-5p | 1.97 | 3.32 | 2.02e-03 | 3.63e-02 | AGAGCUUAGCUGAUUGGUGAAC |
| hsa-miR-92b-5p | 1.95 | 4.26 | 7.85e-04 | 2.22e-02 | AGGGACGGGACGCGGUGCAGUG |
| hsa-miR-629-5p | 1.89 | 4.45 | 6.11e-03 | 6.06e-02 | UGGGUUUACGUUGGGAGAACU |
| hsa-miR-3615 | 1.88 | 2.16 | 1.34e-03 | 2.88e-02 | UCUCUCGGCUCCUCGCGGCUC |
| hsa-miR-941 | 1.86 | 3.04 | 3.85e-04 | 1.48e-02 | CACCCGGCUGUGUGCACAUGUGC |
| hsa-miR-491-5p | 1.77 | 2.52 | 6.75e-04 | 2.05e-02 | AGUGGGGAACCCUUCCAUGAGG |
| hsa-miR-21-5p | 1.76 | 4.44 | 2.66e-02 | 1.30e-01 | UAGCUUAUCAGACUGAUGUUGA |
| hsa-miR-140-3p | 1.76 | 5.02 | 9.78e-04 | 2.52e-02 | UACCACAGGGUAGAACCACGG |
| hsa-miR-629-3p | 1.69 | 2.86 | 4.67e-04 | 1.57e-02 | GUUCUCCCAACGUAAGCCCAGC |
| hsa-miR-151b | 1.69 | 6.32 | 4.26e-07 | 4.56e-05 | UCGAGGAGCUCACAGUCU |
| hsa-miR-193a-3p | 1.61 | 2.51 | 1.71e-02 | 9.98e-02 | AACUGGCCUACAAAGUCCCAGU |
| hsa-miR-484 | 1.60 | 2.39 | 2.43e-03 | 3.81e-02 | UCAGGCUCAGUCCCCUCCCGAU |
| hsa-miR-192-5p | 1.56 | 2.80 | 3.41e-05 | 2.28e-03 | CUGACCUAUGAAUUGACAGCC |
| hsa-miR-21-3p | 1.51 | 4.97 | 7.93e-03 | 6.26e-02 | CAACACCAGUCGAUGGGCUGU |
| hsa-miR-92b-3p | 1.49 | 7.08 | 2.81e-07 | 3.94e-05 | UAUUGCACUCGUCCCGGCCUCC |
| hsa-miR-146b-5p | 1.42 | 2.05 | 1.50e-02 | 9.32e-02 | UGAGAACUGAAUUCCAUAGGCU |
| hsa-miR-625-5p | 1.39 | 4.24 | 2.45e-02 | 1.23e-01 | AGGGGGAAAGUUCUAUAGUCC |
| hsa-miR-505-5p | 1.37 | 2.72 | 7.10e-03 | 6.24e-02 | GGGAGCCAGGAAGUAUUGAUGU |
| hsa-miR-10a-5p | 1.36 | 2.75 | 1.09e-02 | 7.40e-02 | UACCCUGUAGAUCCGAAUUUGUG |
| hsa-miR-3135b | 1.34 | 4.28 | 4.51e-02 | 1.83e-01 | GGCUGGAGCGAGUGCAGUGGUG |
| hsa-miR-2110 | 1.34 | 3.20 | 2.94e-02 | 1.36e-01 | UUGGGGAAACGGCCGCUGAGUG |
| hsa-miR-342-5p | 1.32 | 2.39 | 7.79e-03 | 6.26e-02 | AGGGGUGCUAUCUGUGAUUGA |
| hsa-miR-1307-3p | 1.29 | 7.08 | 2.41e-05 | 1.85e-03 | ACUCGGCGUGGCGUCGGUCGUG |
| hsa-miR-128-3p | 1.29 | 3.65 | 2.03e-02 | 1.12e-01 | UCACAGUGAACCGGUCUCUUU |
| hsa-miR-671-3p | 1.29 | 3.35 | 7.58e-03 | 6.26e-02 | UCCGGUUCUCAGGGCUCCACC |
| hsa-miR-151a-3p | 1.23 | 7.58 | 1.21e-07 | 3.23e-05 | CUAGACUGAAGCUCCUUGAGG |
| hsa-miR-148b-3p | 1.22 | 2.00 | 1.21e-03 | 2.83e-02 | UCAGUGCAUCACAGAACUUUGU |
| hsa-miR-151a-5p | 1.19 | 9.27 | 7.88e-09 | 4.22e-06 | UCGAGGAGCUCACAGUCUAGU |
| hsa-miR-330-3p | 1.18 | 5.02 | 3.20e-02 | 1.44e-01 | GCAAAGCACACGGCCUGCAGAGA |
| ***hsa-miR-10b-5p*** | ***1.16*** | ***2.31*** | ***3.97e-02*** | ***1.69e-01*** | ***UACCCUGUAGAACCGAAUUUGUG*** |
| hsa-miR-3682-3p | 1.14 | 2.04 | 4.80e-03 | 5.25e-02 | UGAUGAUACAGGUGGAGGUAG |
| hsa-miR-425-3p | 1.08 | 3.48 | 1.60e-02 | 9.53e-02 | AUCGGGAAUGUCGUGUCCGCCC |
| hsa-miR-421 | 1.06 | 4.60 | 4.41e-02 | 1.81e-01 | AUCAACAGACAUUAAUUGGGCGC |
| hsa-miR-135b-3p | 1.00 | 2.17 | 1.95e-02 | 1.11e-01 | AUGUAGGGCUAAAAGCCAUGGG |
| hsa-miR-4758-5p | -1.00 | 4.81 | 3.71e-03 | 4.42e-02 | GUGAGUGGGAGCCGGUGGGGCUG |
| hsa-miR-4497 | -1.01 | 10.04 | 6.71e-03 | 6.24e-02 | CUCCGGGACGGCUGGGC |
| hsa-miR-409-3p | -1.05 | 2.12 | 3.74e-02 | 1.61e-01 | GAAUGUUGCUCGGUGAACCCCU |
| hsa-miR-1343-5p | -1.10 | 5.00 | 7.95e-03 | 6.26e-02 | UGGGGAGCGGCCCCCGGGUGGG |
| hsa-miR-548a-3p | -1.11 | 2.80 | 1.67e-02 | 9.83e-02 | CAAAACUGGCAAUUACUUUUGC |
| hsa-miR-193b-3p | -1.14 | 8.74 | 3.26e-06 | 2.91e-04 | AACUGGCCCUCAAAGUCCCGCU |
| hsa-miR-3621 | -1.18 | 5.41 | 8.34e-03 | 6.39e-02 | CGCGGGUCGGGGUCUGCAGG |
| hsa-miR-6126 | -1.36 | 5.86 | 7.60e-03 | 6.26e-02 | GUGAAGGCCCGGCGGAGA |
| hsa-miR-6068 | -1.39 | 3.89 | 2.66e-03 | 3.85e-02 | CCUGCGAGUCUCCGGCGGUGG |
| hsa-miR-503-5p | -1.60 | 7.55 | 4.67e-05 | 2.78e-03 | UAGCAGCGGGAACAGUUCUGCAG |
| hsa-miR-4532 | -1.63 | 6.12 | 3.60e-03 | 4.39e-02 | CCCCGGGGAGCCCGGCG |
| hsa-miR-181b-5p | -1.72 | 8.06 | 2.36e-04 | 1.15e-02 | AACAUUCAUUGCUGUCGGUGGGU |
| hsa-miR-181a-5p | -1.88 | 8.19 | 2.68e-04 | 1.19e-02 | AACAUUCAACGCUGUCGGUGAGU |
| hsa-miR-424-3p | -1.91 | 7.80 | 4.16e-04 | 1.49e-02 | CAAAACGUGAGGCGCUGCUAU |

| | | | | | |
|---|---|---|---|---|---|
| hsa-miR-503-3p | -2.04 | 3.11 | 1.40e-03 | 2.88e-02 | GGGGUAUUGUUUCCGCUGCCAGG |
| hsa-miR-203a | -2.49 | 6.53 | 5.26e-03 | 5.64e-02 | GUGAAAUGUUUAGGACCACUAG |
| hsa-miR-708-5p | -3.50 | 7.93 | 9.89e-04 | 2.52e-02 | AAGGAGCUUACAAUCUAGCUGGG |

Microarray data was RMA background corrected & normalized, SVA batch corrected, foldchange, p-value and FDR were determined by limma package in R.

**Supplementary Table S6: miR-10b target selection and validation.**

| gene | mRNA | P (KM) | log2 (FC) | p-value | r (miR-10) | miR Tarbase miR-10b | Target Scan miR-10 |
|---|---|---|---|---|---|---|---|
| ***DIAPH2*** | **diaphanous-related formin 2 (DIAPH2), transcript variant 156** | **0.02** | **-3.43** | **8.30e-04** | **-0.57** | **TRUE** | **FALSE** |
| *RNF180* | ring finger protein 180 (RNF180), transcript variant 1 | 0.12 | -2.12 | 2.60e-07 | -0.72 | FALSE | TRUE |
| *EPN3* | epsin 3 (EPN3) | 0.22 | -1.54 | 1.39e-03 | -0.70 | FALSE | TRUE |
| *ZNF43* | zinc finger protein 43 (ZNF43), transcript variant 1 | 0.26 | -1.86 | 6.50e-04 | -0.77 | FALSE | TRUE |
| *RBPMS* | RNA binding protein with multiple splicing (RBPMS), transcript variant 1 | 0.34 | -2.46 | 2.60e-05 | -0.78 | FALSE | TRUE |
| *NT5DC3* | 5-nucleotidase domain containing 3 (NT5DC3) | 0.34 | -1.02 | 9.80e-05 | -0.81 | FALSE | TRUE |
| *ZNF470* | zinc finger protein 470 (ZNF470) | 0.38 | -1.13 | 2.10e-06 | -0.78 | FALSE | TRUE |
| *ZFP82* | ZFP82 zinc finger protein (ZFP82) | 0.51 | -1.07 | 1.65e-03 | -0.72 | FALSE | TRUE |
| *RORA* | RAR-related orphan receptor A (RORA), transcript variant 3 | 0.52 | -1.17 | 3.52e-03 | -0.69 | TRUE | TRUE |
| *ARHGEF9* | Cdc42 guanine nucleotide exchange factor (GEF) 9 (ARHGEF9), transcript variant 2 | 0.56 | -1.17 | 8.24e-03 | -0.64 | FALSE | TRUE |
| *SDC1* | syndecan 1 (SDC1), transcript variant 1 | 0.70 | -1.27 | 3.90e-06 | -0.70 | TRUE | TRUE |
| *GJA5* | gap junction protein, alpha 5, 40kDa (GJA5), transcript variant A | 0.73 | -1.30 | 4.07e-03 | -0.67 | FALSE | TRUE |
| *SPARC* | secreted protein, acidic, cysteine-rich (osteonectin) (SPARC), transcript variant 2 | 0.81 | -3.00 | 7.64e-03 | -0.67 | FALSE | FALSE |
| *SH3BGRL* | SH3 domain binding glutamate-rich protein like (SH3BGRL) | 0.85 | -2.33 | 1.30e-06 | -0.68 | FALSE | TRUE |
| *MAP9* | microtubule-associated protein 9 (MAP9) | 0.87 | -2.56 | 7.30e-04 | -0.82 | FALSE | TRUE |
| *SDK2* | sidekick cell adhesion molecule 2 (SDK2) | 0.93 | -1.19 | 4.00e-06 | -0.88 | FALSE | TRUE |
| *ALDH4A1* | aldehyde dehydrogenase 4 family, member A1 (ALDH4A1), transcript variant 3 | 0.95 | -1.02 | 5.10e-06 | -0.76 | FALSE | TRUE |
| *ADAMTS1* | ADAM metallopeptidase with thrombospondin type 1 motif, 1 (ADAMTS1) | 0.96 | -2.98 | 6.10e-05 | -0.74 | FALSE | TRUE |
| *RAB3B* | RAB3B, member RAS oncogene family (RAB3B) | 0.97 | -2.40 | 9.00e-05 | -0.83 | FALSE | TRUE |
| *KAZN* | kazrin, periplakin interacting protein (KAZN), transcript variant D | NA | -1.11 | 8.10e-06 | -0.78 | FALSE | TRUE |

Top 20 *in silico* predicted miR-10 targets, ranked by level of significance (log rank Kaplan-Meier p values, column “p(KM)”) of differences in survival curves when comparing high and mid/low TCGA RNA-seq gene expression datasets of metastatic HNSCC samples. Columns “log2(FC)” and “p-value” indicate foldchange and significance between gene expression of RDEB-SCC and RDEB-KC based on the microarray transcriptome data. Pearson correlation (r) between mean of miR-10a/b log2-transformed normalized microarray expression signals with normalized log2-transformed transcriptome data. Columns “miRTarbase” and “TargetScan” indicate whether the respective the mRNA has previously been reported as miR-10a or miR-10b target (TRUE or FALSE).

**Supplementary Figures:**

**Supplementary Figure S1: miR-10b target identification and selection.** (A) Heat map of significantly deregulated (≥ 2-fold; p ≤ 0.05) miRNAs. Normalized (RMA), batch corrected (SVA) and z-transformed log2 microarray signals were row-wise clustered by single linkage euclidean distance (* indicates miR-10b-5p). (B) KEGG pathway (pwy) enrichment analysis of reported (miRTarbase v6.1) putative targets for ≥ 2-fold, significantly (p-value < 0.05, FDR ≤ 0.2) dysregulated miRNAs (grey dot: RDEB-SCC vs. RDEB-KC, black dot: HC-cSCC vs HC-KC). (C,D) TaqMan PCR (n = 3) on primary mir-10a and -10b transcripts, respectively, confirmed differential regulation of miR-expression. (E) TaqMan PCR (n = 3) on mature miR-10a-5p indicates only mild up-regulation in RDEB-SCC across tested samples. (F) Both, miR-10a and miR-10b were found to be upregulated in microarray analysis. (G) NGS of AGO2-bound miRNAs isolated from immortalized cell lines, comparing DESeq2 normalized read counts of RDEB-SCC to immortalized RDEB-KC, confirmed TaqMan results of consistent upregulation of miR-10b. Error bars represent SEM (C,D,E). Statistical tests: t-test (A,C,D,E) and gene set expression analysis (GSEA) test (B), p < 0.05 was considered as significant.

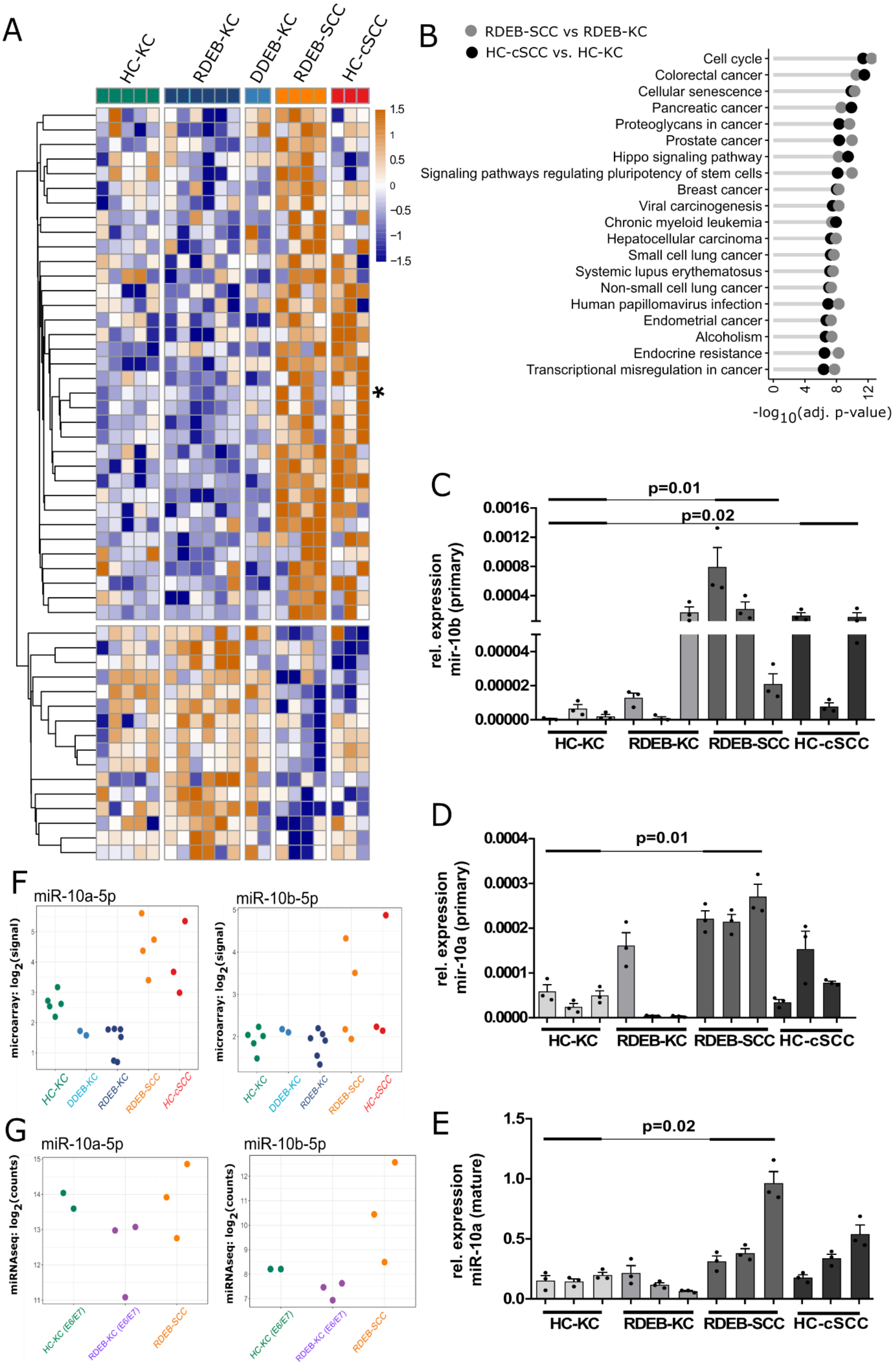
A
HC-KC
RDEB-KC
DDEB-KC
RDEB-SCC
HC-cSCC
1.5
1
0.5
0
−0.5
−1
−1.5
*
B
RDEB-SCC vs RDEB-KC
HC-cSCC vs. HC-KC
Cell cycle
Colorectal cancer
Cellular senescence
Pancreatic cancer
Proteoglycans in cancer
Prostate cancer
Hippo signaling pathway
Signaling pathways regulating pluripotency of stem cells
Breast cancer
Viral carcinogenesis
Chronic myeloid leukemia
Hepatocellular carcinoma
Small cell lung cancer
Systemic lupus erythematosus
Non-small cell lung cancer
Human papillomavirus infection
Endometrial cancer
Alcoholism
Endocrine resistance
Transcriptional misregulation in cancer
0
4
8
12
-log10(adj. p-value)
C
p=0.01
p=0.02
rel. expression mir-10b (primary)
0.0016
0.0012
0.0008
0.0004
0.00004
0.00002
0.00000
HC-KC
RDEB-KC
RDEB-SCC
HC-cSCC
D
p=0.01
rel. expression mir-10a (primary)
0.0004
0.0003
0.0002
0.0001
0.0000
HC-KC
RDEB-KC
RDEB-SCC
HC-cSCC
E
p=0.02
rel. expression miR-10a (mature)
1.5
1.0
0.5
0.0
HC-KC
RDEB-KC
RDEB-SCC
HC-cSCC
F
miR-10a-5p
miR-10b-5p
microarray: log2(signal)
HC-KC
DDEB-KC
RDEB-KC
RDEB-SCC
HC-cSCC
G
miR-10a-5p
miR-10b-5p
miRNAseq: log2(counts)
HC-KC (E6/E7)
RDEB-KC (E6/E7)
RDEB-SCC

296

297

**Supplementary Figure S2: Controls for miRNA probe specificity in *in situ* hybridization on FFPE tissue sections.** (A) H&E staining confirms malignant skin transformation, as opposed to HC-skin. (B) Specificity of miR-10b-5p signal (green) was assessed in tumor tissue by comparison of scrambled negative control probe and $2^{nd}$ step control (no probe) under the same hybridization conditions. Sample processing and hybridization conditions were tested with a positive control probe against *U6* in HC-skin. DAPI was used for nuclei and panKRT (red) for basal KC/SCC staining (C). (D) CD31 and CD45 expression in tissue sections. Immunofluorescence microscopy on HC- and RDEB-SCC tissue sections shows the presence of both, endothelial cells ($CD31^+$) and lymphocytes ($CD45^+$) to different degrees. (E) TWIST1 expression in RDEB-SCC, as shown by immunofluorescence microscopy, and (F) Western blot analysis. TWIST1 is highly expressed in RDEB-SCC cell cultures, as compared to HC-KCs and RDEB-KCs. Alpha-actinin (ACTA) was used as loading control. In immunofluorescence microscopy, TWIST1 was found to be expressed in RDEB-SCC tumor sections, but not in HC-skin.

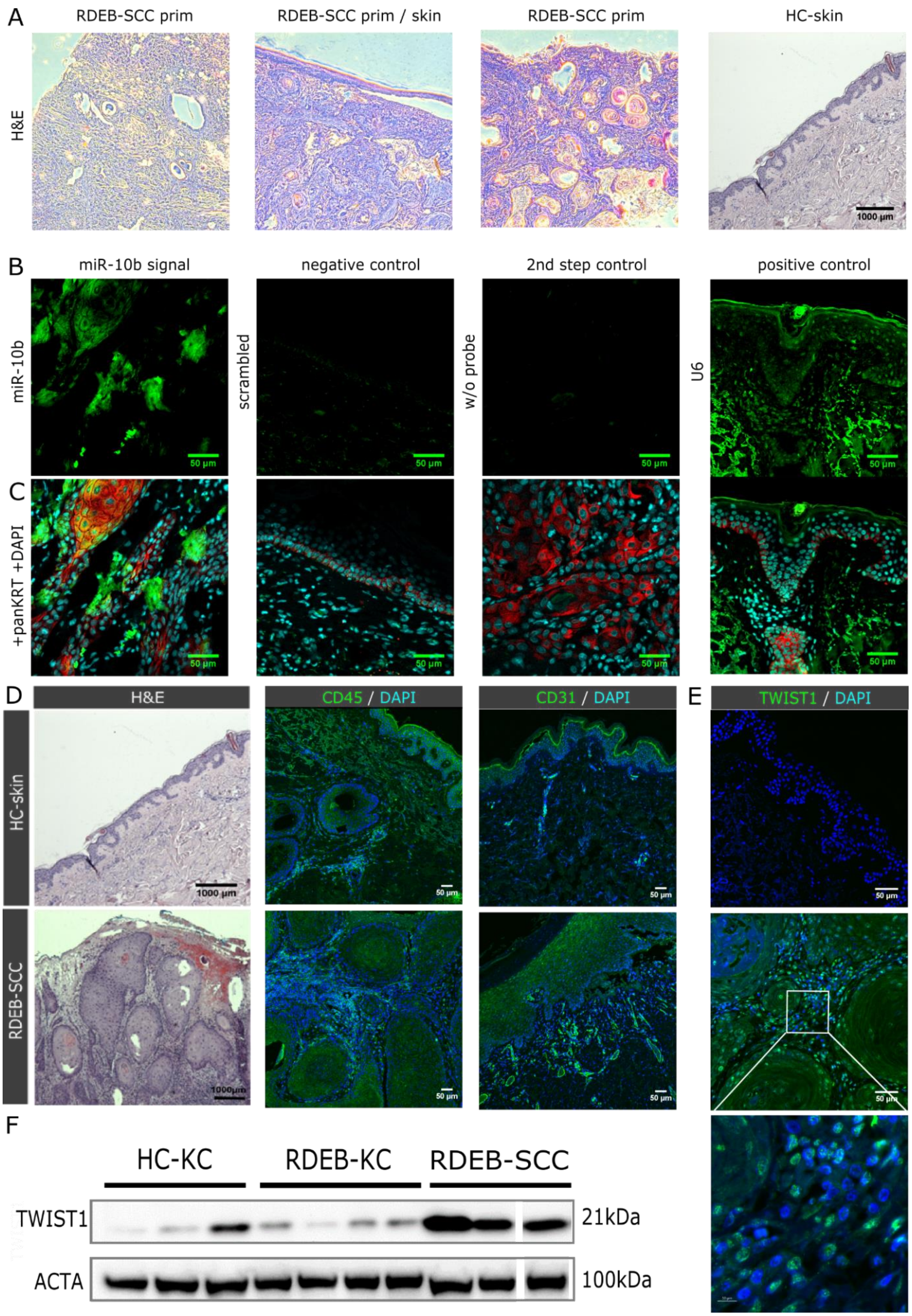
A
RDEB-SCC prim
RDEB-SCC prim / skin
RDEB-SCC prim
HC-skin
H&E
1000 µm
B
miR-10b signal
negative control
2nd step control
positive control
miR-10b
scrambled
w/o probe
U6
50 µm
C
+panKRT +DAPI
D
H&E
CD45 / DAPI
CD31 / DAPI
E
TWIST1 / DAPI
HC-skin
RDEB-SCC
F
HC-KC
RDEB-KC
RDEB-SCC
TWIST1
21kDa
ACTA
100kDa

**Supplementary Figure 3: miRNA probe specificity for *in situ* hybridization on cultured cells and single cell resolution image-based data analysis workflow.** Images were converted to grey scale using the CellProfiler software and the miR-10b specific signal (green) was used for segmentation of cytoplasm (cellular mask) and DAPI (blue) for nuclei (nuclear mask) (A,B). miR-10b specific signals were cumulated within cellular masks for each single cell and reported as "integrated intensity" after subtraction of cumulated nuclear signal, which was most likely derived from unspecific DNA-binding (C,D). Scale bar = 50 µm.

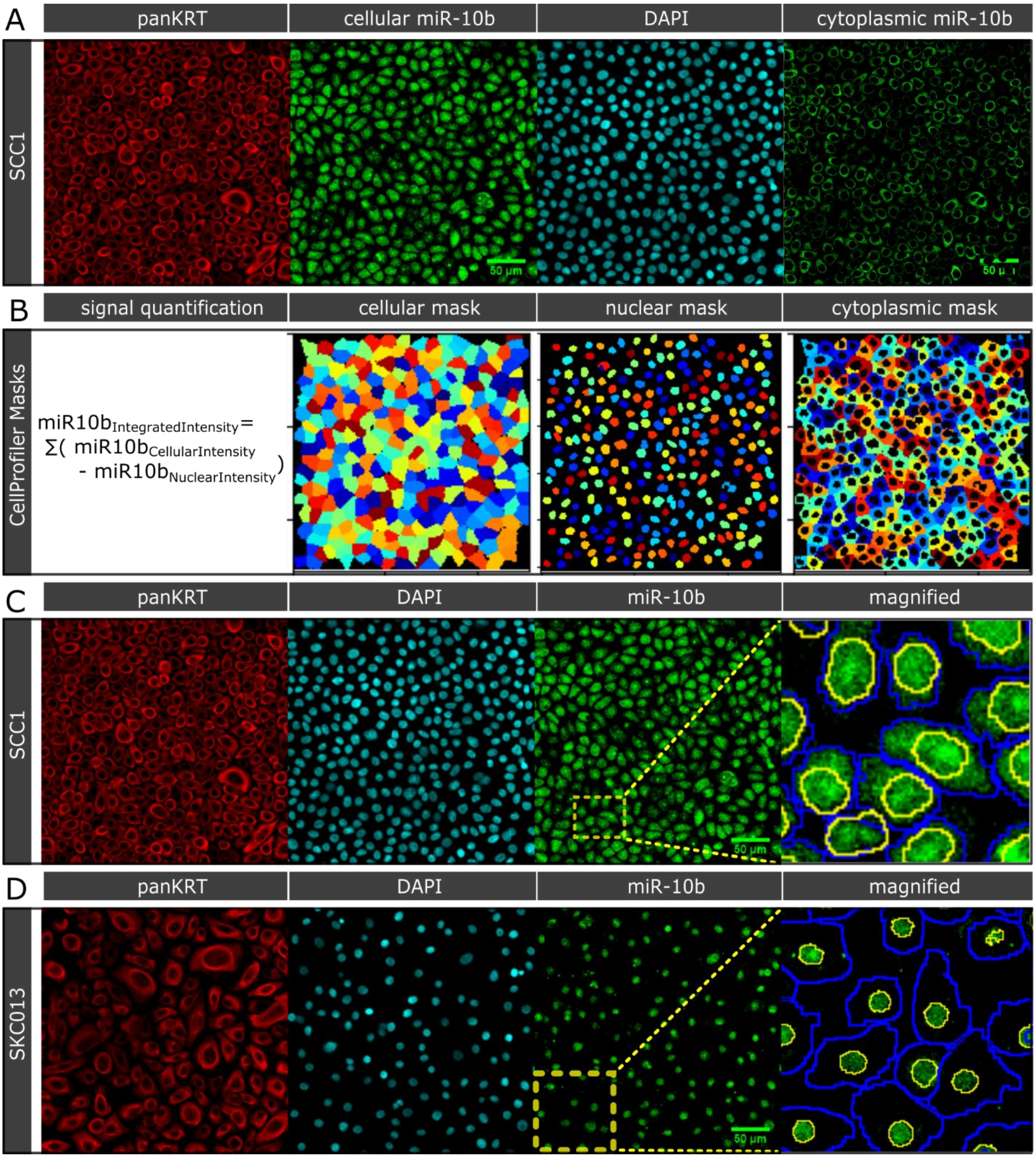

**Supplementary Figure S4: miRNA probe specificity for *in situ* hybridization on cultured cells.** Panel (A) demonstrates miR-10b-5p signal (FITC) in RDEB-SCC1 cells. DAPI was used for nuclei and panKRT (red) for cytoplasmic basal KC/SCC staining. (B) Absence of signal when applying a scrambled negative control probe at same hybridization conditions as used for miR-10b-5p probe is shown. In addition a second step control, without any probe was performed to verify non-specific staining of anti-DIG Fab-FITC (C). (D) A probe for *U6* (green) was used as positive control. Scale bar = 50 µm.

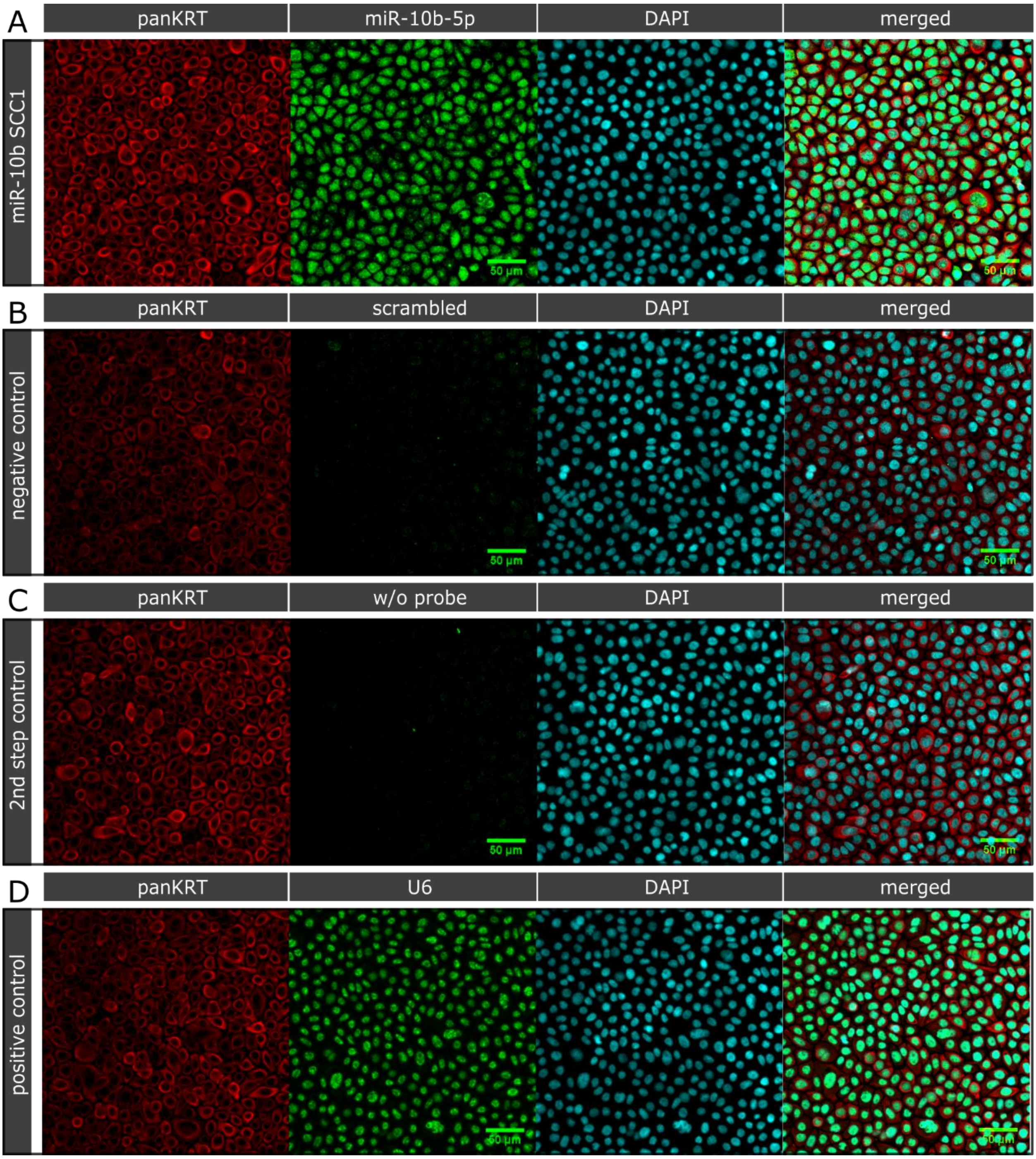

**Supplementary Figure S5: Generation of experimental cell lines.** (A) Two different non-malignant keratinocyte lines were retrovirally transduced with a miR-10b expression cassette under the control of an *U6* promoter. Expression levels of mature miR-10b were assessed by TaqMan PCR (n = 3; error bars represent SEM). (B) sqRT-PCR confirmed reduced levels of *DIAPH2* in miR-10b overexpressing keratinocytes (n = 3; error bars represent SEM). (C) Upon transfection of immortalized 1090KC with *DIAPH2* targeted CRISPR/Cas9, single clones were analyzed for successful disruption of *DIAPH2*. Two clones harboring a homozygous deletion of one nucleotide within exon 1 were used for further experiments. (D) Western blot analysis confirmed loss of *DIAPH2* expression upon CRISPR-mediated disruption. (E,F) $1090KC^{DIAPH2-/-}$ cells showed a delayed gap closure, similar to miR-10b overexpressing cells (* $p < 0.05$; $n \geq 8$; t-test; error bars represent SEM). The percentage (%) of gap closure compared to the starting area at 0 hrs is given at 3, 6, 9 and 12 hrs. (G,H) While 1090KC barely form stable aggregates *in vitro*, knock-out of the miR-10b target *DIAPH2* conferred the capacity to form stable aggregates. Size distribution of aggregates is similar to that of RDEB-SCC cells. Representative images of aggregates in parental 1090KC compared to $1090KC^{DIAPH2-/-}$ are shown. (I) Viability of cells in aggregates was confirmed via live (calcein-AM) – dead (ethidium) staining. (J,K) CRISPR/Cas9 knock-out of MIR10B in RDEB-SCC1 was confirmed by Sanger sequencing, as well as TaqMan PCR on bulk population and clones generated by one round of serial dilution (* $p < 0.05$; n = 3; unpaired t-test; error bars represent SEM).

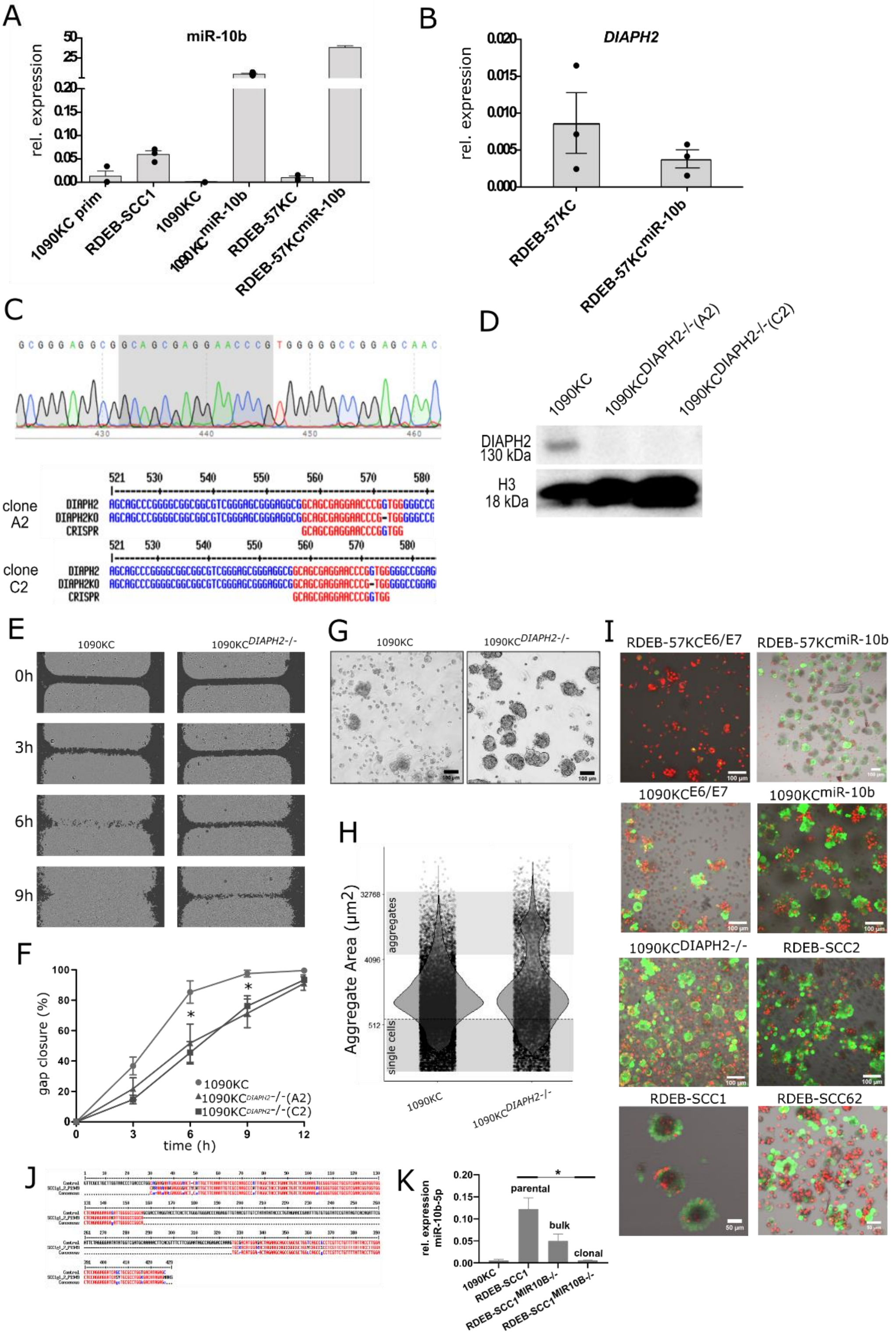
A
miR-10b
rel. expression
1090KC prim
RDEB-SCC1
1090KC
1090KC miR-10b
RDEB-57KC
RDEB-57KC miR-10b
B
DIAPH2
rel. expression
RDEB-57KC
RDEB-57KC miR-10b
C
clone A2
clone C2
DIAPH2
DIAPH2KO
CRISPR
D
1090KC
1090KC DIAPH2-/-(A2)
1090KC DIAPH2-/-(C2)
DIAPH2 130 kDa
H3 18 kDa
E
1090KC
1090KC DIAPH2-/-
0h
3h
6h
9h
F
gap closure (%)
time (h)
1090KC
1090KC DIAPH2-/-(A2)
1090KC DIAPH2-/-(C2)
G
1090KC
1090KC DIAPH2-/-
H
Aggregate Area (µm2)
aggregates
single cells
1090KC
1090KC DIAPH2-/-
I
RDEB-57KC E6/E7
RDEB-57KC miR-10b
1090KC E6/E7
1090KC miR-10b
1090KC DIAPH2-/-
RDEB-SCC2
RDEB-SCC1
RDEB-SCC62
J
K
rel. expression miR-10b-5p
parental
bulk
clonal
1090KC
RDEB-SCC1
RDEB-SCC1 MIR10B-/-
RDEB-SCC1 MIR10B-/-

**Supplementary Figure S6: Characterization of experimental cell lines.** (A) Generation of cell lines and morphologic characteristics. Experimental cells had different sizes, as evaluated by measuring the median cell surface area. For each cell type ≥ 10 cells were measured. Those cells stably expressing miR-10b were significantly smaller in size than their parental cells. Also upon *DIAPH2* disruption, cells became slightly smaller. Analysis of CD44 and CD24: (B) RDEB-57KC overexpressing miR-10b and respective parental cells were analyzed by flow cytometry for surface proteins CD44 and CD24. In the presence of miR-10b, the ratio of $CD44^{high}$ / $CD24^{-/low}$ increased from 2.82% to 11.37%. Both cell lines expressed similar levels of CD44. (C) RDEB-SCC cell lines showed overall high expression levels of CD44 and different subpopulations of CD24 expressing cells.

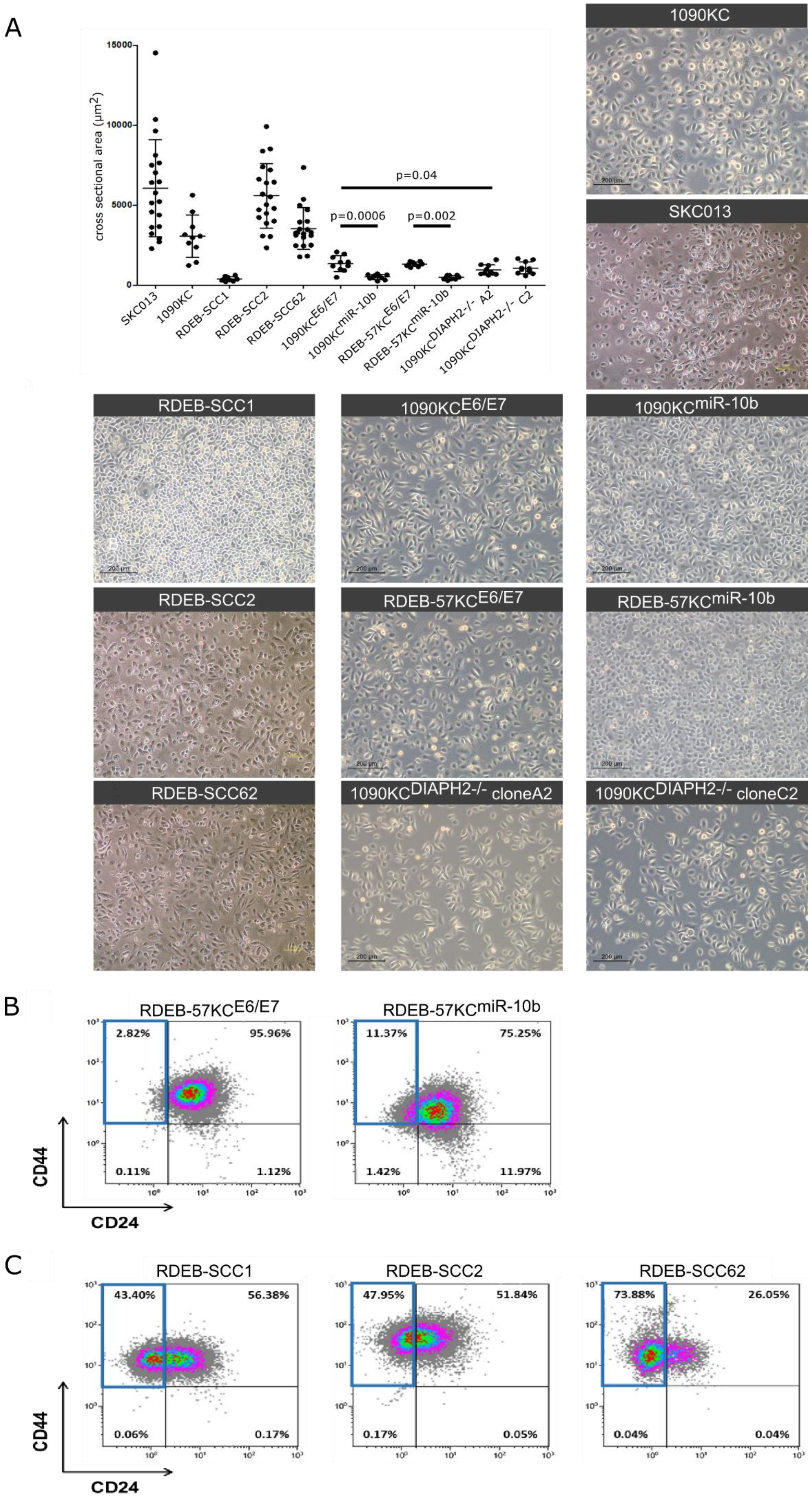
A
15000
10000
5000
0
cross sectional area (μm²)
p=0.04
p=0.0006
p=0.002
SKC013
1090KC
RDEB-SCC1
RDEB-SCC2
RDEB-SCC62
1090KCE6/E7
1090KCmiR-10b
RDEB-57KCE6/E7
RDEB-57KCmiR-10b
1090KCDIAPH2-/- A2
1090KCDIAPH2-/- C2
1090KC
SKC013
RDEB-SCC1
1090KCE6/E7
1090KCmiR-10b
RDEB-SCC2
RDEB-57KCE6/E7
RDEB-57KCmiR-10b
RDEB-SCC62
1090KCDIAPH2-/- cloneA2
1090KCDIAPH2-/- cloneC2
B
RDEB-57KCE6/E7
2.82%
95.96%
0.11%
1.12%
RDEB-57KCmiR-10b
11.37%
75.25%
1.42%
11.97%
CD44
CD24
C
RDEB-SCC1
43.40%
56.38%
0.06%
0.17%
RDEB-SCC2
47.95%
51.84%
0.17%
0.05%
RDEB-SCC62
73.88%
26.05%
0.04%
0.04%
CD44
CD24

**Supplementary Figure S7: Validation of miR-10b targets HOXD10 and DIAPH2.** (A) Scientific literature (1,000 Pubmed-listed articles) was screened for terms “cancer” and “miR-10b” on which named entity recognition (NER) was performed to extract biological relevant terms (geniatagger). A generated word cloud highlights prominent terms, among others, HOXD10. (B) Validation of *HOXD10* expression by sqRT-PCR did not reveal a downregulation of *HOXD10* mRNA in RDEB-SCC and HC-cSCC lines compared to controls (n = 3; error bars represent SEM; t-test; dashed bars: normalized to GAPDH, plain bars: normalized to TUBA). However, at the protein level HOXD10 was downregulated as shown by immunofluorescence microscopy (C) on cells (HOXD10: red) and (D) on tissue sections of, RDEB-skin versus an RDEB-SCC and HC skin versus HC-cSCC (HOXD10: green, DAPI: blue). (E) Mean microarray log2 expression signals of miR-10a and miR-10b correlate well (r = 0.77; Pearson) with ΔCq values of miR-10b TaqMan qPCR. Normalized (RMA) log2-transformed microarray expression signals of ≥ 2-fold deregulated genes (RDEB-SCC vs. RDEB-KC) of the transciptome panel for miR-10 targets in either miRTarbase or Targetscan, were correlated with mean of normalized (RMA) log-2 transformed miR-10 sample-matched microarray expression signals. Among those, DIAPH2 showed a significant ($p < 0.01$) correlation (r = - 0.57; Pearson). (F) A dual luciferase assay confirmed the direct interaction of a miR-10b mimic with the *HOXD10* 3’UTR and (G) the *DIAPH2* 3’UTR, both cloned downstream the luciferase gene. Luciferase signals decreased when co-transfected with miR-10b mimic on average by 37% for *HOXD10* and 50% for *DIAPH2*, respectively, compared to a scrambled control (n = 6; error bars represent SEM; t-test). (H) Upon transfection of HC-KCs with 50 mM miR-10b mimic, DIAPH2 protein levels decreased compared to cells transfected with scrambled control in dosimetric Western blot analysis (n = 5; normalized to annexin 1; error bars represent SEM; t-test) (I) Validation of *DIAPH2* expression by sqRT-PCR showed downregulation of *DIAPH2* mRNA in RDEB-SCC compared to HC-KC (n = 3; error bars represent SEM; t-test; dashed bars: normalized to GAPDH, plain bars: normalized to TUBA). (J) DIAPH2 expression levels were lower in RDEB-SCC1 compared to HC-KC, whereas levels were increased in a bulk population of RDEB-SCC1$^{MIR10B-/-}$. Representative images from two indepentent repeats (DIAPH2: green, DAPI: blue).

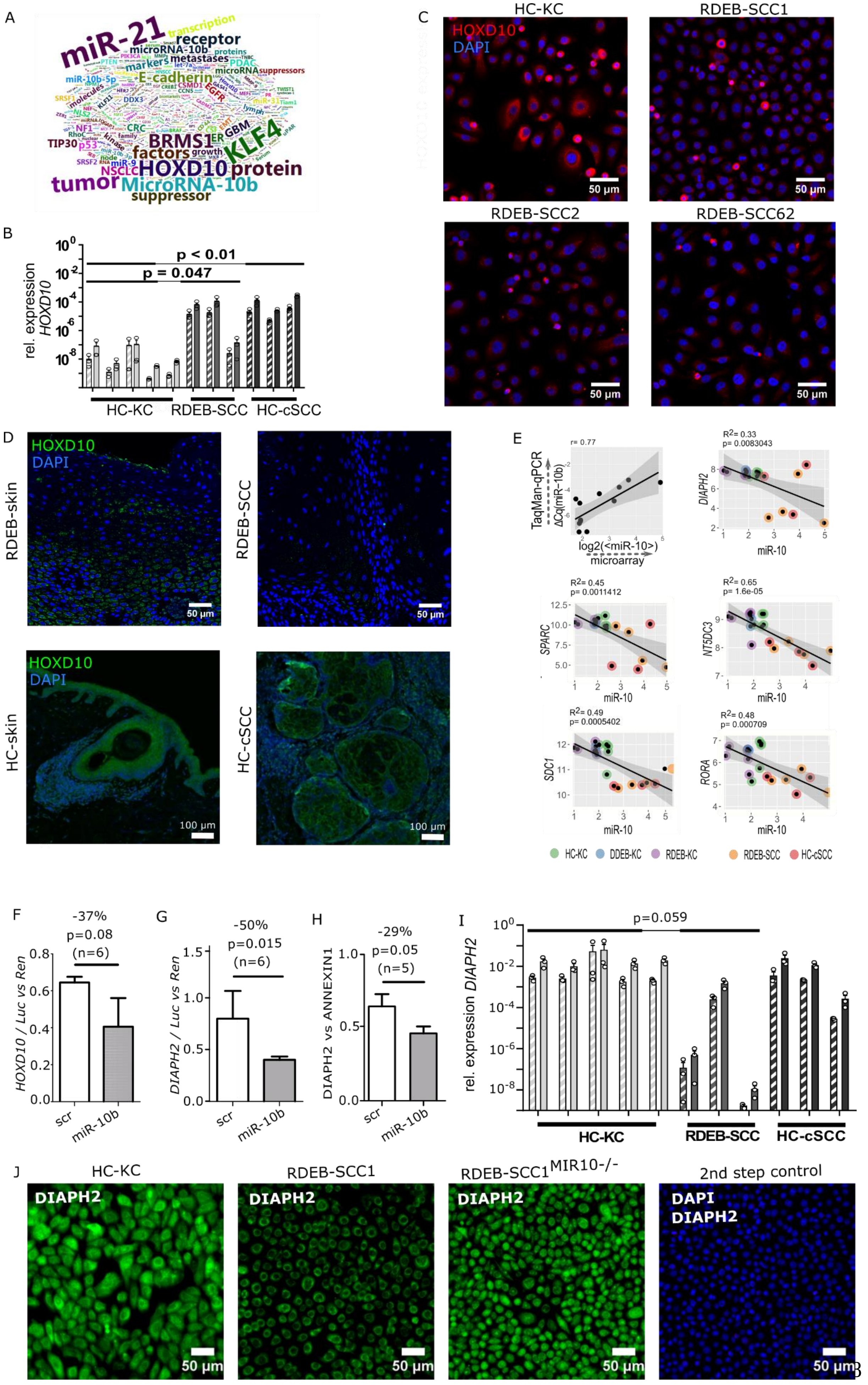
A
miR-21
HOXD10 protein
MicroRNA-10b
tumor
KLF4
BRMS1
E-cadherin
B
rel. expression HOXD10
p < 0.01
p = 0.047
HC-KC
RDEB-SCC
HC-cSCC
C
HC-KC
RDEB-SCC1
RDEB-SCC2
RDEB-SCC62
HOXD10
DAPI
50 µm
D
HOXD10
DAPI
RDEB-skin
RDEB-SCC
HC-skin
HC-cSCC
50 µm
100 µm
E
r= 0.77
TaqMan-qPCR
ΔCq(miR-10b)
log2(<miR-10>)
microarray
R2= 0.33
p= 0.0083043
DIAPH2
miR-10
R2= 0.45
p= 0.0011412
SPARC
R2= 0.65
p= 1.6e-05
NT5DC3
R2= 0.49
p= 0.0005402
SDC1
R2= 0.48
p= 0.000709
RORA
HC-KC
DDEB-KC
RDEB-KC
RDEB-SCC
HC-cSCC
F
-37%
p=0.08
(n=6)
HOXD10 / Luc vs Ren
scr
miR-10b
G
-50%
p=0.015
(n=6)
DIAPH2 / Luc vs Ren
H
-29%
p=0.05
(n=5)
DIAPH2 vs ANNEXIN1
I
p=0.059
rel. expression DIAPH2
HC-KC
RDEB-SCC
HC-cSCC
J
HC-KC
RDEB-SCC1
RDEB-SCC1 MIR10-/-
2nd step control
DIAPH2
DAPI
50 µm

**Supplementary Figure S8: Survival analysis and target evaluation.** (A) Survival analysis of miR-10 targets in TCGA HNSCC RNAseq datasets. Stratification of patients based on miR-10 target expression levels results in *DIAPH2* as most significant impact factor on survival (Low: n = 64, high: n = 22, log-rank test). (B) miR-10b targets *DIAPH2*, but not *DIAPH1 and 3,* shows lower expression in RDEB-SCC (n = 8) compared to RDEB-skin (n = 10; Wilcox test) tissue. (C) As expected, *HOXD10* expression is not reduced at the mRNA level, as it is regulated by translational inhibition. Normalized RNA-seq data generated by Cho *et al* (20), data was retrieved from GEO repository (GSE111582).

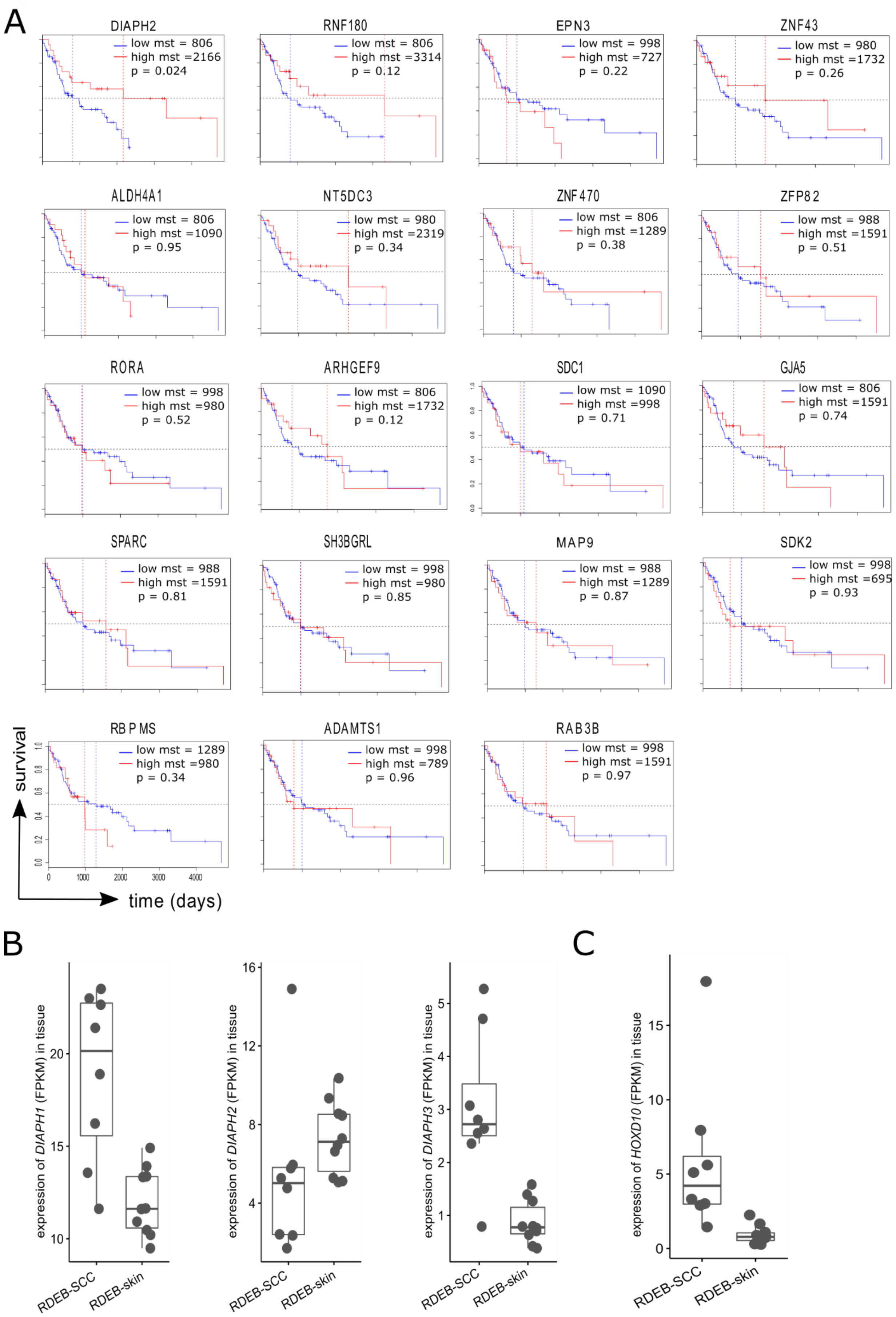
A
DIAPH2
low mst = 806
high mst =2166
p = 0.024
RNF180
low mst = 806
high mst =3314
p = 0.12
EPN3
low mst = 998
high mst =727
p = 0.22
ZNF43
low mst = 980
high mst =1732
p = 0.26
ALDH4A1
low mst = 806
high mst =1090
p = 0.95
NT5DC3
low mst = 980
high mst =2319
p = 0.34
ZNF470
low mst = 806
high mst =1289
p = 0.38
ZFP82
low mst = 988
high mst =1591
p = 0.51
RORA
low mst = 998
high mst =980
p = 0.52
ARHGEF9
low mst = 806
high mst =1732
p = 0.12
SDC1
low mst = 1090
high mst =998
p = 0.71
GJA5
low mst = 806
high mst =1591
p = 0.74
SPARC
low mst = 988
high mst =1591
p = 0.81
SH3BGRL
low mst = 998
high mst =980
p = 0.85
MAP9
low mst = 988
high mst =1289
p = 0.87
SDK2
low mst = 998
high mst =695
p = 0.93
RBPMS
low mst = 1289
high mst =980
p = 0.34
ADAMTS1
low mst = 998
high mst =789
p = 0.96
RAB3B
low mst = 998
high mst =1591
p = 0.97
survival
time (days)
B
expression of DIAPH1 (FPKM) in tissue
expression of DIAPH2 (FPKM) in tissue
expression of DIAPH3 (FPKM) in tissue
RDEB-SCC
RDEB-skin
C
expression of HOXD10 (FPKM) in tissue
